# Single-cell and spatial transcriptomics unveils key regulators governing cell differentiation for *Schistosoma japonicum* sexual development

**DOI:** 10.64898/2026.01.20.700708

**Authors:** Zhigang Lu, Xiaoxu Wang, Shun Li, Xuxin Li, Pengfei Du, Yameng Zheng, Lu Liu, Yanyan Zhang, Omri Wurtzel, Guofeng Cheng

**Affiliations:** Shanghai Tenth People’s Hospital, Clinical Center for Brain and Spinal Cord Research, Affiliated Shanghai Blue Cross Brain hospital, Institute for Infectious Diseases and Vaccine Development, Tongji University School of Medicine, 500 Zhen-nan Road, Shanghai 200331, China; Faculty of Natural Sciences, University of Hohenheim, Stuttgart, 70599, Germany; School of Biotechnology, Jiangsu University of Science and Technology, Zhen Jiang, 212100, China; Shanghai Veterinary Research Institute, Chinese Academy of Agricultural Sciences, Key Laboratory of Animal Parasitology of Ministry of Agriculture and Rural Affairs, Shanghai, 200241, China; The School of Neurobiology, Biochemistry & Biophysics, George S. Wise Faculty of Life Sciences, Tel Aviv University, Tel Aviv, Israel

**Keywords:** *Schistosoma japonicum*, Single-cell RNA sequencing, Spatial transcriptomics, Transcription factors, Development, Sexual maturation

## Abstract

Schistosomiasis remains a major global health challenge, primarily due to the large number of egg production by sexually mature parasites. However, the cellular basis and regulatory programs governing sexual development and egg production are poorly characterized. Here, we constructed a dynamic single-cell atlas of *Schistosoma japonicum* (*S. japonicum*) covering key development stages of sexual maturation and egg production, and integrated it for the first time with spatial transcriptomics to map tissue–resolved cellular niches. Through bioinformatic analysis and experimental validation, we identified several critical cell populations and regulatory networks that drive parasite development and egg laying. By mapping *de novo* identified transcription factors (TFs) onto these atlases, we generated a comprehensive spatiotemporal expression profile of TFs and define the regulatory programs of several tissue-specific TFs including *Zfp*, *Fbp3*, and *Lim*. Integrated re–clustering further revealed and validated key regulators of germline stem-cell differentiation in *S. japonicum*. Finally, comparative analyses with *S. mansoni* single-cell datasets revealed both conserved and species–divergent cellular plasticity, providing unprecedented insights into their distinct biological traits. Collectively, our comprehensive study establishes a foundational resource for understanding schistosome biology, deciphers regulatory programs underlying cell differentiation and pathogenicity, and accelerates the identification of key molecules involved in sexual development against schistosomiasis.

## 1 Introduction

Schistosomiasis is a devastating neglected tropical disease that affects over 250 million people worldwide^1^. The lack of an effective vaccine and the continued reliance on a single drug, praziquantel (PZQ), highlight the urgent need to identify novel targets and develop new strategies for disease control ^2^. A deeper understanding of the key biological events underlying parasite pathogenicity is therefore particularly important for discovering effective targets.

Schistosomes are dioecious flatworms whose development is tightly regulated ^1^. It had been recognized for a long time that male-female pairing is essential for gamete production, sexual maturation of the female, and egg production^3–6^, which is a primary driver of host pathology and disease transmission. Defining the cellular basis and differentiation programs involved in these events could reveal key target nodes for reducing pathology and blocking transmission. While *Schistosoma* cell populations have been analyzed by single-cell RNA-seq (scRNA-seq), these studies focused on the discrete stages/organs—including miracidia^7^, sporocysts^8^, schistosomula^9^, and adults^10,11^ —as well as for specific reproductive organs^12,13^, the dynamic cellular shifts from juvenile to sexually mature adult worms remain uncharacterized, especially in *S. japonicum*. This critical developmental window is directly responsible for egg production and subsequent disease pathology, yet its underlying regulatory landscape is largely unknown.

Here, we used scRNA-seq to systematically profile dynamic cell populations across three critical developmental stages of *Schistosoma japonicum* (*S. japonicum*): pre-pairing schistosomula (16 days post-infection, D16), paired adults (D20), and sexually mature adults (D26). By generating six single-cell atlases spanning these stages and integrating them for the first time with spatial transcriptomics, we mapped tissue niches and defined cellular localizations. We then conducted genome-wide identification of transcription factors (TFs) and combined with single-cell data, constructed a comprehensive spatiotemporal map of TF dynamics. Several TF-driven programs were validated experimentally and shown to be essential for sexual maturation and egg production. Integrated re-clustering further identified and validated key regulators of germline stem-cell (GSC) differentiation. Moreover, cross–species comparison with *S. mansoni* revealed both conserved and species–divergent cellular features, providing mechanistic insights into their distinct biological traits. These results including scRNA-seq atlases from multiple stages and spatial transcriptomics across almost annotated tissues, and species-specific cell transcriptomes have been organized as a publicly available, interactive online resource (https://schisto.xyz/sj-atlas/). Collectively, our comprehensive study provides a foundational molecular and cellular framework for understanding schistosome biology, reveals key regulators of pathogenicity-linked cell differentiation, and highlights potential targets against schistosomiasis.

## 2 Results

### 2.1 *Schistosoma japonicum* scRNA-seq and spatial transcriptomic atlases

Using an optimized dissociation protocol and the 10 × Genomics Chromium platform, we obtained 127,411 single-cell transcriptomes from six libraries (**Figure 1A**). After stringent quality control, 60,029 high-quality cell transcriptomes in the combined six libraries were retained and clustered into 62 transcriptionally distinct clusters (denoted C#, where # indicates cluster identity, **Figure 1B** and Supplementary **Table 1**, showing a list of cluster marker genes in each cell cluster). These clusters included neurons (10 clusters), muscles (8), tegument and tegument progenitors (7 and 4), neoblasts and their progeny (5 and 9), flame cells (3), parenchyma (3), and multiple reproductive populations—including S1 and S1 progeny, vitellocytes (early/late/mature), germline stem cells (GSCs) and their progeny, late–stage germ cells from the female ovary (named as “Female late”) and male testis (named as “Male late”), Mehlis gland cells, and two ambiguous clusters (Supplementary **Table 1**). These 62 clusters were further consolidated into 21 biologically annotated cell types, representing nearly all known schistosome tissues (Supplementary **Figure 1**). Whole-mount in situ hybridization (WISH) using the probes for representative marker genes confirmed the anatomical localizations of these cell identities in adult worms (**Figure 1C** and Supplementary **Figure 1**). To spatially resolve cell architecture, we performed Stereo–seq spatial transcriptomics on sections of adult *S. japonicum*. Our optimized protocol generated high–resolution maps comprising 14,323 spatially barcoded cells. These spatial maps faithfully recapitulated all cell clusters from the single–cell atlas **(Figure 1D**, Supplementary **Table 2** and Supplementary **Figure 2A and 2B**) and resolved their anatomical distributions. Notably, they captured spatial sections of male and female gonads (**Figure 1D** and Supplementary **Figure 2B**) as well as the male–female pairing interface (Supplementary **Figure 2B** and Supplementary **Table 3**).

**Figure 1.**
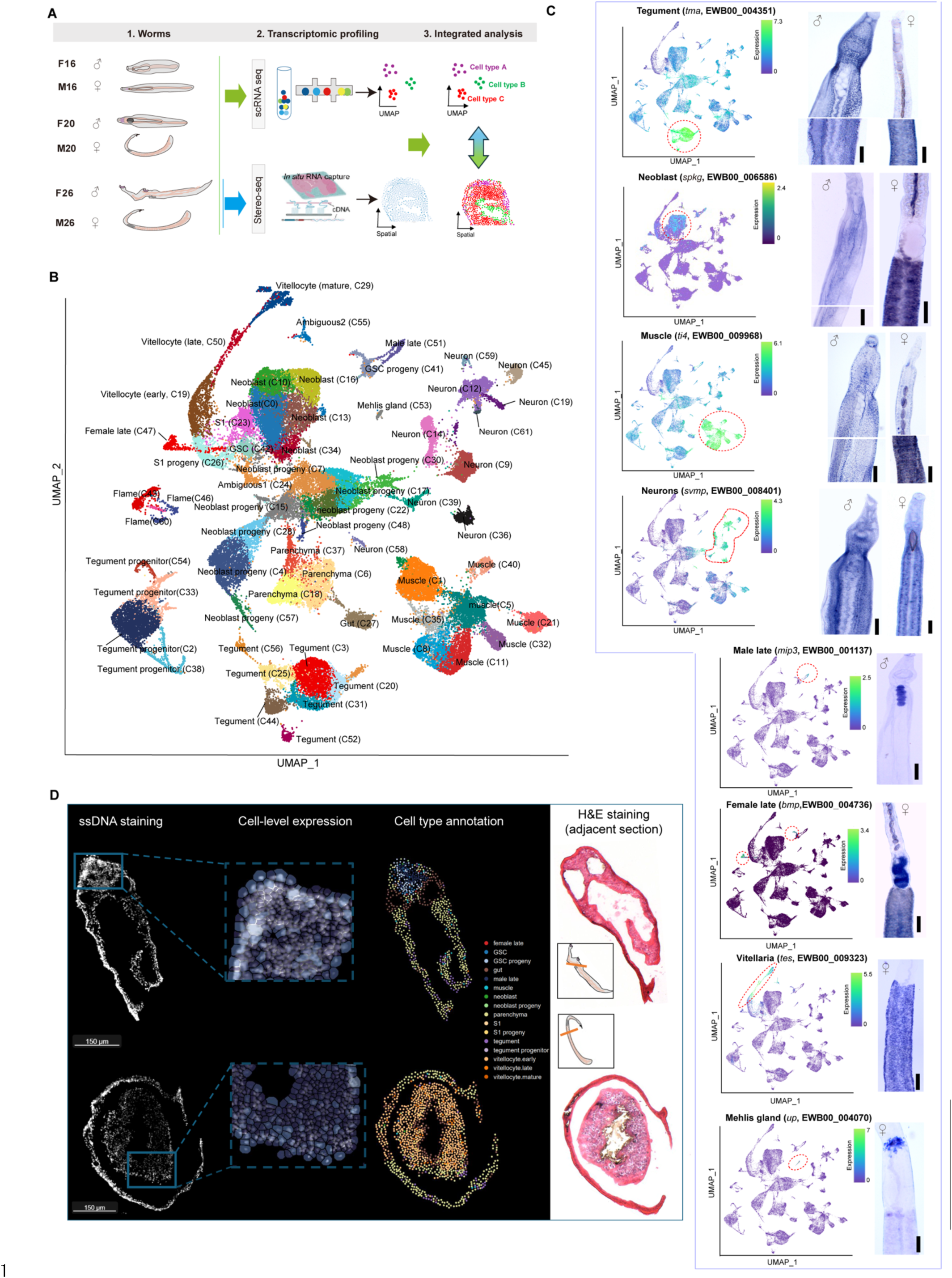
*S. japonicum* single-cell and spatial transcriptomic atlases. **(A)** Schematic overview of scRNA-seq and spatial transcriptomics analyses for *S. japonicum*. **(B)** UMAP plot of 62 transcriptionally distinct cell clusters identified through scRNA-seq. **(C)** Validation of anatomical localizations of representatively identified tissue cells. *tma* (Tegument membrane-associated antigen, EWB00_004351), *skpg* (Serine/threonine-protein kinase greatwall, EWB00_006586), *ti4* (Tropomyosin-2 isoform 4E, WB00_009968), *svmp* (Synaptic vesicle membrane protein VAT-1 isoform 1, EWB00_008401), *mip3* (M-phase inducer phosphatase 3, EWB00_001137), *bmp* (Bone marrow proteoglycan, EWB00_004736); *tes* (Trematode Eggshell Synthesis, EWB00_009323), *up* (Uncharacterized protein EWB00_004070). Red dashed lines on UMAPs highlight the relevant cell cluster. Data are representative results of 20-50 investigated worms. Scale bars = 200 µm. **(D)** Spatial transcriptomic mapping of cell types within anatomical structures of male and female sections. Cells in the spatial transcriptomics were identified based on the ssDNA staining image.

Our single–cell atlas revealed extensive somatic heterogeneity in *S. japonicum* (**Figure 1B**), aligning with reported complexity in *S. mansoni*^11^. Neuronal diversity is particularly prominent, comprising ten transcriptionally distinct clusters. While all clusters highly express the synaptic vesicle gene *Vat-1* (EWB00_008401) (Supplementary **Figure 3**), each exhibits a unique transcriptional signature, reflecting a compartmentalized nervous system comprising central ganglia and peripheral networks^14^. Double-fluorescence *in situ* hybridization (FISH) confirmed the spatial segregation of neuronal subpopulations marked by co-expression of *Vat-1* with either a tyrosine kinase (C45) or a calcium-uptake protein (C59) (Supplementary **Figure 3C**). Muscle tissue similarly displayed considerable heterogeneity, forming eight distinct clusters defined by high expression of *Actin* (EWB00_002502), *Troponin I* (EWB00_011315), and *Smoothelin* (EWB00_002930) (Supplementary **Figure 4**). These clusters express conserved developmental regulators, including highly active SMAD/BMP, Wnt, and Notch signaling pathways, highlighting the evolutionary conservation of myogenic programs. The tegument, a critical interface for nutrient uptake, parasite survival, and host immune modulation, also showed substantial diversity, with seven transcriptionally discrete clusters identified (Supplementary **Figure 5**). Parenchymal cells, which fulfil essential structural and metabolic roles, are resolved into three clusters (Supplementary **Figure 6**). Together, these results provide a high-resolution, spatially annotated transcriptomic atlas for *S. japonicum*, providing an integrative framework for systematically investigating cell functions, intercellular crosstalk, and molecular programs that underpin schistosome development and reproductive maturation.

### 2.2 Cellular landscapes across *S. japonicum* sexual development

The stage- and sex-specific morphological divergence observed in *Schistosoma* is underpinned by distinct cellular composition^15^. To reveal the dynamic of cell composition during *S. japonicum* sexual development, we comparatively analyzed these six single-cell atlases (**Figure 2A**). Notably, females and males showed broadly similar cellular profiles at D16 and D20 (**Figure 2A**) including parenchyma, muscle, neurons, tegument, neoblasts, neoblast progeny and tegument progenitors (**Figure 2B and 2C**). At these stages, neoblasts (11-24% of total cells), neoblast progeny (18-26%), muscle (10-26%), neurons (8.6-17%), and tegument (8.5-18%) constituted the most abundant lineages, underscoring their critical roles in supporting schistosomula development (**Figure 2B and 2C**).

**Figure 2.**
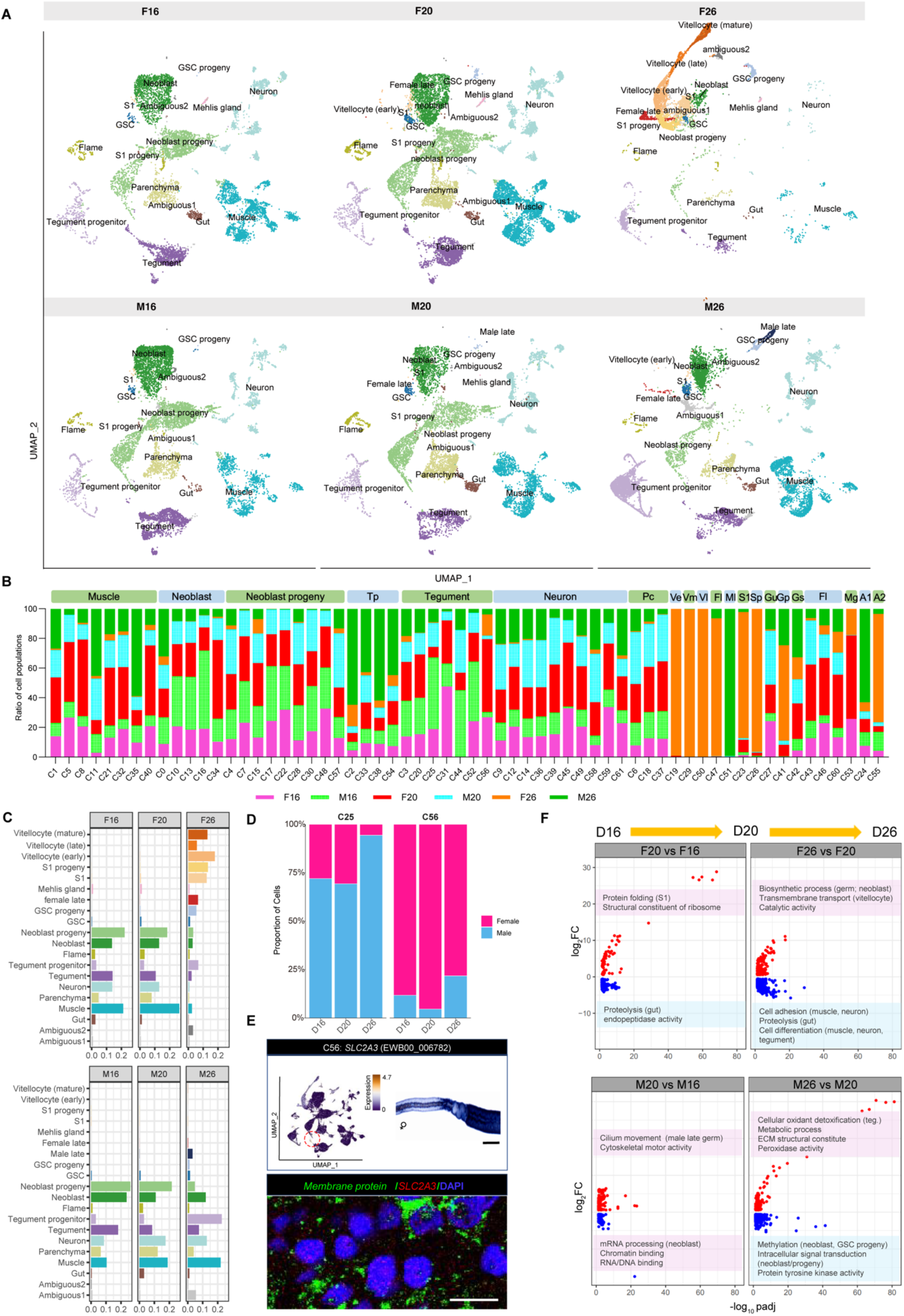
Landscapes of scRNA-seq atlases and dynamic cell populations during *S. japonicum* sexual development. **(A)** UMAP plot of scRNA-seq atlases from male and female worms across different stages**. (B)** Percentage of each cell cluster for three stages of males and females. Muscle (C1, C5, C8, C11, C21,C32, C35, and C40); Neoblast (C0, C10, C13, C16, C34); Neoblast progeny (C4, C7, C15, C17, C22, C28, C30, C48, C57); Tp, tegument progenitor (C2, C33, C38, C54); Tegument (C3, C20, C25, C31, C44, C52, C56); Neuron (C9, C12, C14, C36, C39, C45, C49, C58, C59, C61); Pc, parenchyma (C6, C18, C37); Ve, vitellocyte (early) (C19) ; Vm, vitellocyte (mature) (C29) ; Vl, vitellocyte (late) (C50) ; Fl, female late (C47); Ml, male late (C51);S1 (C23); Sp, S1 progeny (C26); Gu, gut (C27) ; Gp, GSC progeny (C41); Gs, GSC (C42); Fl, flame (C43, C46, C60); Mg, mehlis gland (C53); A1, ambiguous 1 (C24); A2, ambiguous 2 (C55). (**C**) Dynamic cell population shifts during *S. japonicum* sexual development. **(D)** Proportions of female and male cells in C25 and C56 clusters across different stages. **(E)** Spatial localization of the C56 tegumental subcluster, visualized by WISH and double FISH using the marker SLC2A3 (EWB00_006782), indicating enrichment surrounding the ovarian duct and testis regions. Scale bars = 200 µm for WISH and 10 µm for FISH. **(F)** Gene Ontology (GO) analysis of differentially expressed genes from dynamic cell populations across different stage between males and females.

Sex-specific cell types emerged upon male-female pairing at D20, including rare gametes (*n* = 2 cells) and early vitellocytes (C19; *n* = 11 cells), reflecting the coupling between pairing and the onset of gametogenesis (**Figure 2B and 2C**). When parasites reached full sexual maturity (D26), distinct cellular trajectories associated with egg production appeared. In females, gametogenic and vitellogenic lineages as well as S1 and S1 progeny expanded substantially (from 0.78% at D20 to 62.12% at D26). This expansion coincided with a pronounced reduction in muscle (−24.02%), neoblast progeny (−14.99%), and neuronal populations (−12.23%). In contrast, males displayed a marked increase in tegument progenitors (+19.69%) (**Figure 2B and 2C**).

Comparative analyses of cell composition across these stages revealed dynamic shifts in both somatic and stem cell populations. For example, most muscle cell clusters (6 of 8; C1, C5, C8, C21, C31, and C40) were enriched in females at D16 and D20 but shifted toward male-biased enrichment upon sexual maturation at D26 (**Figure 2A and 2B**). Tegument populations also displayed compartmentalized sexual dimorphism: C25 was male-enriched, whereas C56 was female-enriched (**Figure 2D**). Spatial analysis using double FISH by applying the probes of the solute carrier gene *SLC2A3* (EWB00_006782, a marker of C56) localized this population to the oviduct interface (**Figure 2E**). In addition, we identified transient cell populations likely supporting stage-specific functions (**Figure 2B**). For example, C31 was detectable in juveniles but almost absent in adults, suggesting a potential role specific to the schistosomulum stage. Moreover, Mehlis’ gland cells were also detected in pre-paired females (D16), indicating the possible functions that is independent of egg production (**Figure 2B and 2C**).

Further analyses of sequential development scRNA-seq trajectories uncovered markedly divergent functional shifts (**Figure 2F**). In females, development transitioned from an early emphasis on protein folding and ribosome assembly during pairing to programs dominated by germline/neoblast biosynthesis and vitellocyte transport at D26. Concurrently, genes involved in pathways such as proteolysis and cell adhesion were downregulated. In males, pairing was associated with heightened ciliary motility and cytoskeletal activity, followed by the induction—by D26—of oxidoreductase activity, extracellular matrix remodeling, and peroxidase function, whereas chromatin/RNA–binding and intracellular signaling pathways were suppressed. These sex–specific cellular activities provide important clues to the coordinated molecular strategies that allocate metabolic and structural resources to support sustained egg production.

### 2.3 Dynamic cellular transcriptomes across parasite sexual development

To elucidate the molecular basis of cellular dynamics during sexual development, we performed a global analysis of transcriptional changes using our scRNA-seq data. Pseudo–bulk analyses revealed highly concordant transcriptomes between immature males and females (D16, D20), indicating shared somatic and progenitor states at these stages (**Figure 3A** and Supplementary **Figure 7A**). In contrast, mature females (F26) exhibited extensive transcriptional divergence—both from their immature stages (F16, F20) and from mature males (M26)—reflecting large-scale reprogramming associated with sexual maturation (**Figure 3A** and Supplementary **Figure 7A**). Notably, mature males (M26) remained transcriptionally similar to their immature stages (M16 and M20), suggesting the sustained core somatic and progenitor functions throughout the assessed developmental window (**Figure 3A**).

**Figure 3.**
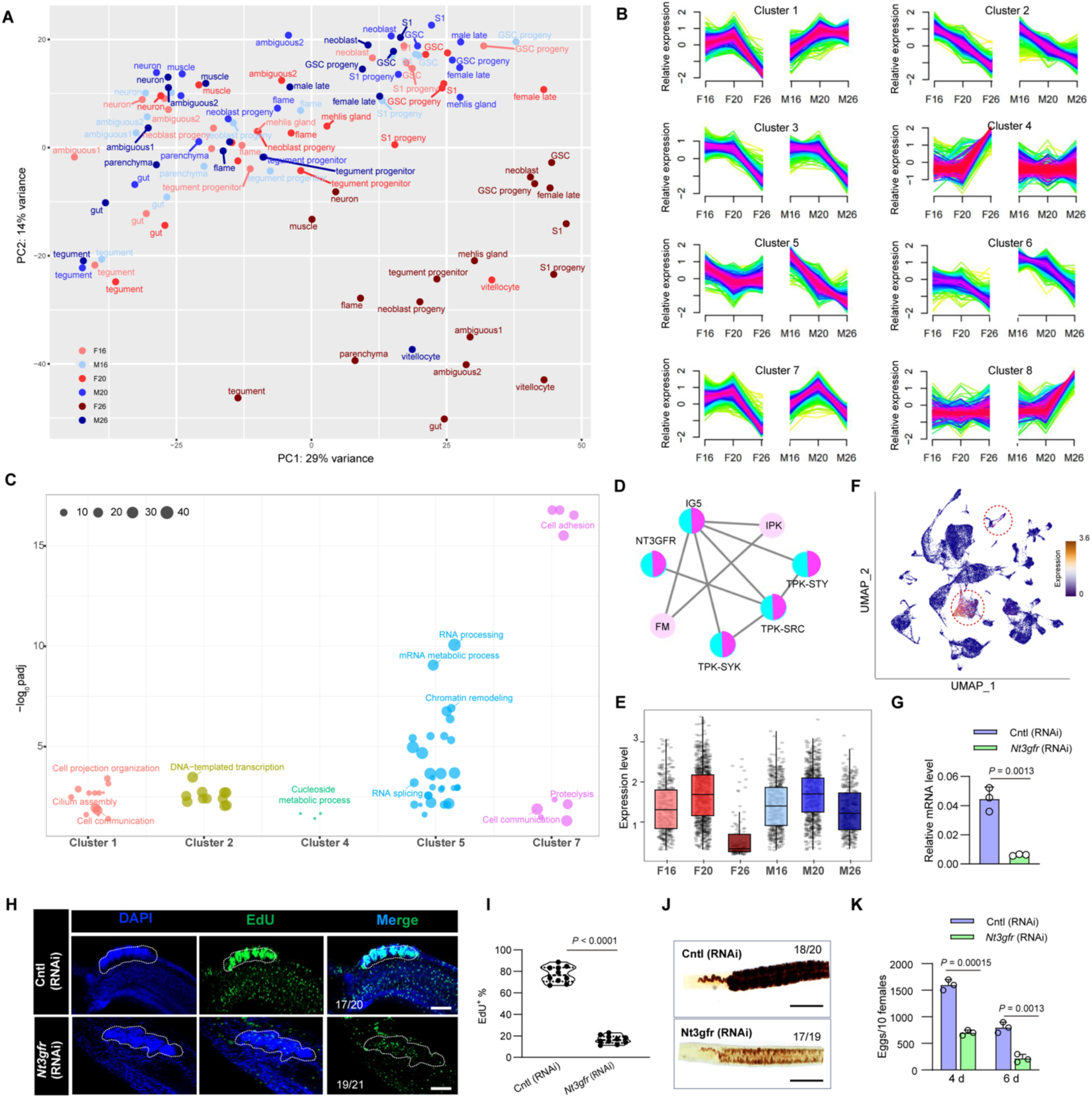
The dynamics of cellular transcriptome during *S. japonicum* sexual maturation. (**A**) PCA analysis of cell types across different stages between males and females. (**B**) Fuzzy clustering of dynamically expressed genes of cell populations across different stages between males and females. (**C**) Enriched biological processes in genes of selected clusters from (b). (**D**) Prediction of protein-protein interaction network for cluster 7 by STRING. IG5 (Integrin alpha-5, EWB00_002919); NT3GFR (NT-3 growth factor receptor, EWB00_000911); IPK (Integrin-linked protein kinase, EWB00_010398); TPK-STY (Tyrosine-protein kinase, EWB00_001471); TPK-SRC (Tyrosine-protein kinase, EWB00_007520); TPK-SYK (Tyrosine-protein kinase, EWB00_011403); FM (Fermitin family 1, EWB00_002103). (**E**) Expression level of *Nt3grf* in non-zero cells in different stages of the scRNA-seq data. (**F**) UMAP plot of *Nt3grf* expression in parenchyma and male late cells. (**G**) RT-qPCR analysis of the transcript level of *Nt3grf* in suppressed males. Parasites were collected 6 days post-treatment for RNA extraction. Data represent mean ± SD from three biological replicates. (**H**) EdU staining analysis of cell proliferation from testis in *Nt3grf* KD males at 6 days of post treatment. The image shown is representative of results from 17-21 worms. Numbers given indicate the fraction of worms that were similar with respect to the phenotype in relation to the total number of worms examined. Scale bars = 200 μm. (**I**) Quantifications of EdU^+^ positive cells from (H) by Image J. n = 10. (**J**) FastBlue BB staining showing morphological vitellaria in the females co-cultured with *Nt3grf* KD males for 6 days. The image shown is representative of results from 17-20 females. Numbers given indicate the fraction of worms that were similar with respect to the phenotype in relation to the total number of worms examined. Scale bars = 400 μm. (**K**) Reduced egg production in the females co-cultured with *Nt3grf* KD males at 4 and 6 days. Data illustrate representative results as mean ± standard error derived from the experiments of triplicates under two biological replicates.

Through fuzzy clustering analyses, we resolved temporal expression patterns of 2,670 dynamic genes into eight clusters (**Figure 3B** and Supplementary **Table 4**). Their cellular origins and putative functions were annotated (**Figure 3C** and Supplementary **Figure 7B**): Genes in cluster 1 (progressively increased in males, peaking in paired males) are primarily from muscle cells with the enrichment for calcium–ion binding, ciliary function, cell adhesion and Wnt signaling; Genes in cluster 2 (downregulated in both sexes) are from tegument and neoblasts that linked to DNA-templated transcription; Genes in cluster 7 (transiently upregulated during pairing) are predominantly from parenchymal cells that associated with cell adhesion, cell communications; Genes in cluster 8 (male–specific genes induced after pairing) are from male late germ cells and flame cells that related to ciliary motility; Genes in cluster 4 are mainly from vitellocytes with transmembrane transport functions.

Given the critical role of males during pairing in assisting female sexual development and egg production^3–6^, we focused on peaking expression genes at this stage, particularly those in males-enriched Clusters 7. STRING analysis of these genes found that NT–3 growth factor receptor (NT3GFR) mediated tyrosine kinase pathway may involve in this process (**Figure 3D**). Additionally, *Nt3gfr* was notably upregulated when comparing cellular transcriptomes of paired worms (D20) and pre-paired worms (D16) (This analysis led to the identification of 63 significantly upregulated and 91 downregulated genes, Supplementary **Table 5**). Analyses of scRNA-seq data found that *Nt3gfr* was high expression in parenchyma and testis of adult males (**Figure 3E and 3F**). RNAi-mediated knockdown (KD) of *Nt3gfr* impaired cell differentiation in both testis and parenchyma (**Figure 3H and 3I**). When co-cultured with untreated females, *Nt3gfr* KD males were shown to influence vitellarium development of females (**Figure 3J**) associated with a significant reduction in egg production (**Figure 3K**). Overall, these results suggest that *Nt3gfr^+^* parenchyma cells in males could be involved in the regulation of vitellarium development and egg production of the female.

### 2.4 Spatiotemporal map of transcription factors during parasite sexual development

To elucidate how transcription factors (TFs) regulate cell differentiation during *Schistosoma* sexual development, we performed a genome–wide identification of TFs in *S. japonicum*, defining a repertoire of 181 TFs (Supplementary **Table 6**). Integration with our scRNA-seq atlas generated a spatiotemporal map of 65 dynamically expressed TFs across development (**Figure 4A**). STRING analysis revealed that 52 of these TFs assemble into three potential regulatory modules centered on RNA polymerase II–dependent transcription, homeobox proteins, and homeobox/HLH hybrids (**Figure 4B**), suggesting that their cooperative interactions could coordinate parasite development and sexual maturation.

**Figure 4.**
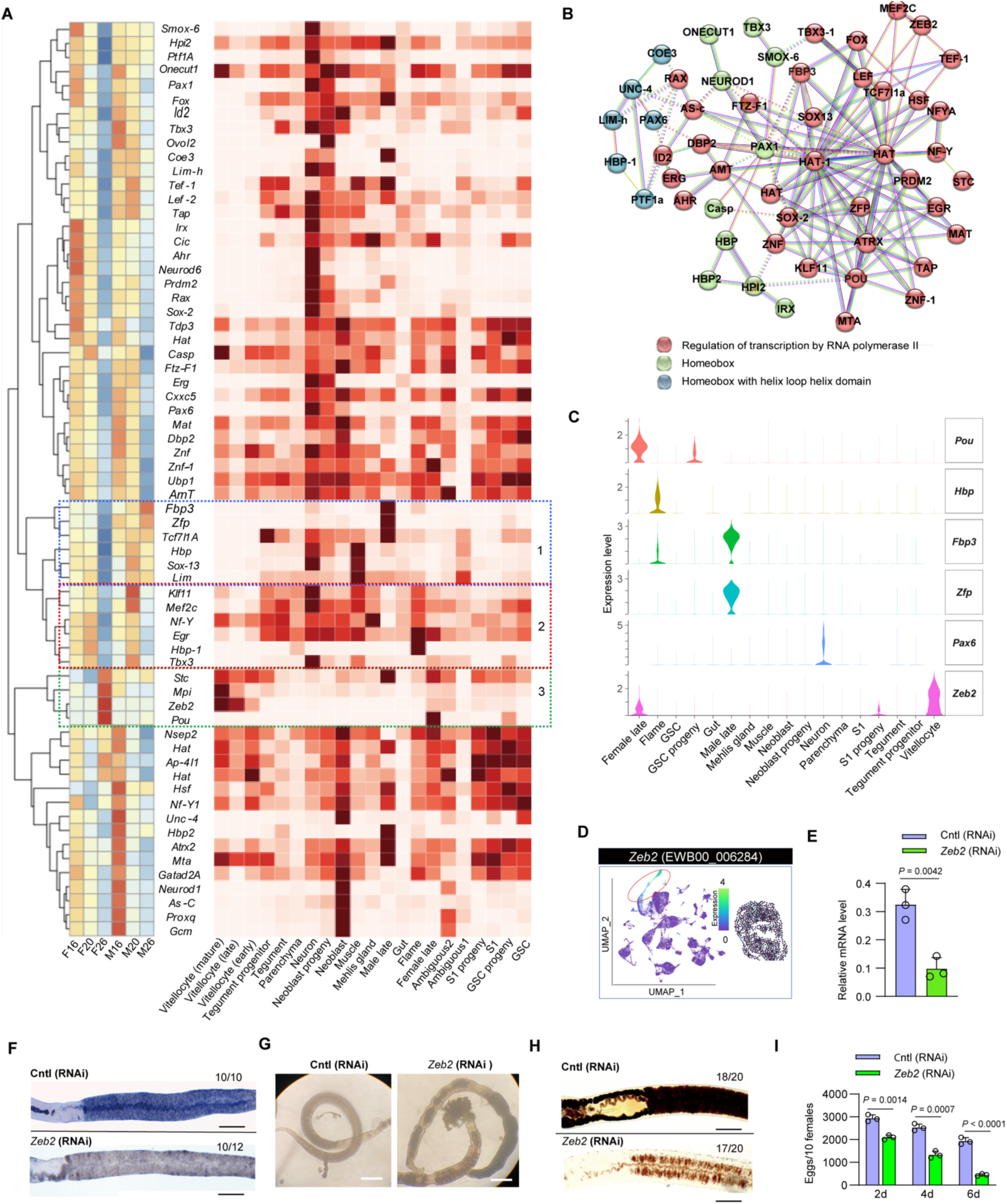
Spatiotemporal mapping of transcription factors (TFs) in identified cell populations during sexual maturation and validation of *Zeb2* function. **(A)** Heatmap showing the dynamic expression profiles of TFs in different cells at different stages. **(B)** Proposed interaction network of identified TFs. (**C**) Violin plots showing the tissue specific expressions of selected TFs. (**D**) UMAP plot and spatial transcriptomics showing *Zeb2* expression in vitellocytes. (**E**) RT-qPCR analysis of the transcript level *of Zeb2* in the inhibited females. Parasites were collected at 6 days of post treatment and then subject to RNA isolation. Data illustrate representative results indicating mean and standard deviation obtained from an experiment conducted in triplicate under two biological replicates. (**F**) WISH analysis of the reductio of *Zeb2* transcript in KD females. A representative image is shown from 10-12 females. Numbers in upper right indicate fraction of worms that are similar to those presented/total number of worms examined. Scale bars = 400 μm. (**G**) Morphological alterations of females treated with *Zeb2* dsRNA. Scale bars = 1000 μm. (**H**) FastBlue BB staining showing morphological vitellaria in *Zeb2* KD females at 6 days post-treatment. The image shown is representative of results from 17-20 females. Numbers given indicate the fraction of worms that were similar with respect to the phenotype in relation to the total number of worms examined. Scale bars = 400 μm. (**I**) *Zeb2* inhibition led to a decreased number of eggs of paired females following RNAi. Data illustrate representative results as mean ± standard error derived from the experiments of triplicates under two biological replicates.

By focusing on TFs exhibiting peak expression around pairing or maturation, we classified 16 TFs into three groups (**Figure 4A**): Group 1 TFs are highly expressed in males after pairing but progressively decreased in females (Blue box in Figure 4A); Group 2 TFs are highly expressed in both sexes during pairing (D20) (Pink box in Figure 4A); and Group 3 TFs are high expression specify in female during egg production (Green box in Figure 4A). Notably, several TFs showed striking cell–type specificity: *Hbp* (*Homeobox protein*, EWB00_011065) in flame cells, *Pax6* (*Paired box protein*, EWB00_004658) in neurons, *Zeb2* (*Zinc finger E-box-binding homeobox protein*, EWB00_006284) in vitellocytes, *Zfp* (*Zinc finger protein*, EWB00_005374) and *Fbp3* (*Forkhead box protein isoform 3*, EWB00_002486) in testis (male late), and *Pou* (*POU domain transcription factor 1*, EWB00_008527) in the ovary (female late) (**Figure 4C**).

Focusing on the vitellaria—a key organ that produces yolk cells for egg assembly—*Zeb2* was shown to be highly expressed in vitellocytes, as confirmed by scRNA-seq results and spatial transcriptomics (**Figure 4D**) as well as WISH (**Figure 4F**, Cntl (RNAi). RNAi–mediated knockdown (KD) of *Zeb2* in females significantly reduced its transcript level, verified by RT–qPCR (**Figure 4E**) and WISH (**Figure 4F**). Notably, we found that *Zeb2* suppression led to pronounced morphological defects (**Figure 4G**), disrupted cell proliferation of vitellocytes (**Figure 4H** and Supplementary **Figure 8**) and markedly decreased egg production (**Figure 4I**). Together, these results indicate that *Zeb2* plays important roles in regulation of cell proliferation in vitellarium and thus potentially contributes to egg production in *S. japonicum*.

### 2.5 Male gonad-enriched *Zfp^+^* and *Fbp3*^+^ cells regulate testis development

While *Zeb2* delineates a female–specific regulatory role in vitellaria, several TFs showed elevated expression in paired males, suggesting their potential roles in sustaining male functions that may support female development and egg production. Spatiotemporal results highlighted two such TFs: *Zfp* (*Zinc finger protein*, EWB00_005374) and *Fbp3* (*Forkhead box protein isoform 3*, EWB00_002486). Both scRNA-seq data and spatial transcriptomics indicated their high expressions in male late germ cells (**Figure 5A**). RNAi–mediated knockdown (KD) of *Zfp* or *Fbp3* in adult males significantly reduced their transcript levels (**Figure 5B and 5C**) and impaired cell activities in testes—an effect more pronounced upon *Zfp* suppression (**Figure 5D and 5E**)—while ovarian proliferation remained unaffected, confirming their male–specific functions (Supplementary **Figure 9**).

**Figure 5.**
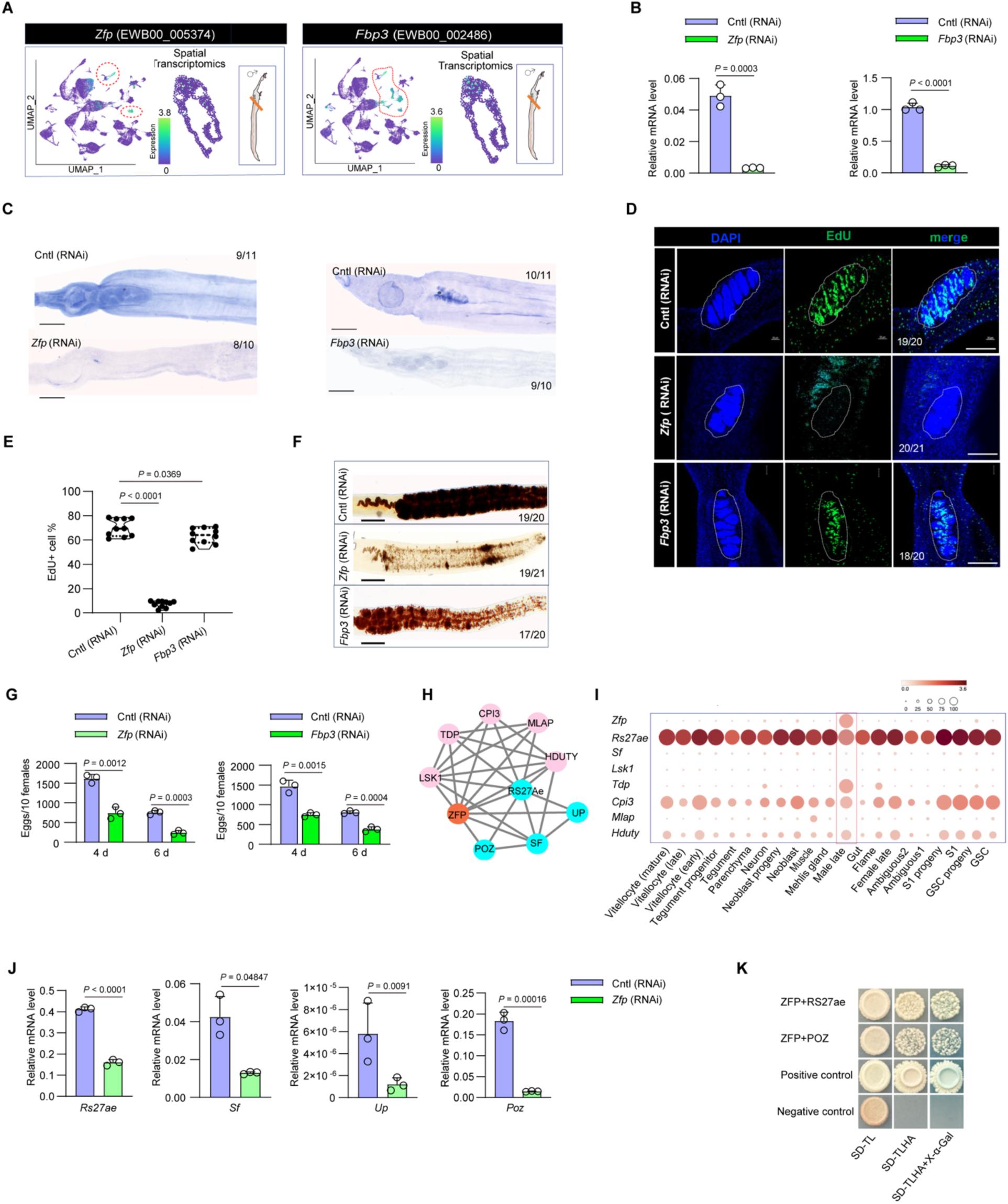
Characterization of *Znf*^+^ and *Fbp3*^+^ cell functions in *S. japonicum* testis. **(A)** UMAP plot and spatial transcriptomic analysis indicated localization of *Znf⁺* and *Fbp3⁺* cells. **(B)** RT-qPCR analysis of the transcript level of *Znf* or *Fbp3* in the suppressed males. Data illustrate representative results indicating mean and standard deviation obtained from an experiment conducted in triplicate. This is a representative result from two biological replicates. (**C**) WISH analysis showing *Zfp⁺* and *Fbp3^+^* cell distribution in control and dsRNA-treated males. A representative image is shown based on data from 8-11 males. Numbers given indicate the fraction of worms that were similar with respect to the phenotype in relation to the total number of worms examined. (**D**) EdU staining analysis of cell proliferation in the testis of *Zfp* and *Fbp* KD males. Shown is a representative image based on 18-21 examined males. Numbers given indicate the fraction of worms that were similar with respect to the phenotype in relation to the total number of worms examined. Scale bars= 200 μm. (**E**) Quantification of EdU^+^ positive cells from (D) by Image J. (**F**) FastBlue BB staining showing vitellaria in female worms following co-culture with *Zfp* and *Fbp* KD males. Female schistosomes were stained with FastBlue BB on days 12 after co-culture with *Zfp* and *Fbp* KD males, and stained worms were imaged using a microscopy. Shown is a representative image from 17-21 males. Numbers given indicate the fraction of worms that were similar with respect to the phenotype in relation to the total number of worms examined. Scale bars = 200 μm. (**G**) Egg production was significantly decreased in females co-cultured with *Zfp* and *Fbp* KD males. Data illustrate representative results as mean ± standard error derived from the experiments of triplicates. (**H**) ZFP mediated protein-protein interaction by STRING analysis. LSK1 (Leucine-rich repeat serine/threonine-protein kinase 1, EWB00_010175); MLAP (Muscle M-line assembly protein, EWB00_009314); SF (Suppressor of fused, EWB00_006102); TDP (Thioredoxin domain-containing protein 3, EWB00_007811); CPI3 (Chromodomain-helicase-DNA-binding protein isoform 3, EWB00_009542); POZ (Speckle-type POZ protein-like B, EWB00_003497); UP (Uncharacterized protein, EWB00_010575); RS27Ae (Putative small subunit ribosomal protein S27Ae, EWB00_000191); HDUTY (Histone demethylase UTY, EWB00_007844). (**I**) ScRNA-seq data analysis of the expressions of putative interactors in male late cells. (**J**) RT-qPCR analysis of the transcript levels of the putative interactor in *Zfp* KD males. Parasites were collected at 6 days of post treatment and then subject to total RNA isolation. Data illustrate representative results indicating mean and standard deviation obtained from an experiment conducted in triplicate under three biological replicates. (**K**) Validation of protein interactions by yeast two hybrid assay.

Then, we determined whether knockdown of *Zfp* or *Fbp3* in males affects egg production in females. After thorough washing, dsRNA–treated males were co–cultured with untreated females. Females co-cultured with *Zfp* or *Fbp3* suppressed males exhibited impaired vitellarium development—more severe in the *Zfp* KD treatment (**Figure 5F**) —and significantly reduced egg production (**Figure 5G**). STRING analysis suggested that ZFP may interact with IFT122, RS27ae, POZ, Sufu and other partners, potentially linking to the Hedgehog signal pathway (**Figure 5H**). ScRNA–seq data further demonstrated that several of these interactors were co–expressed with *Zfp* in late male germ cells (**Figure 5I**). Moreover, RT-qPCR analyses of *Zfp* KD males displayed significant downregulation of *Rs27ae*, *Sufu*, *Usp*, and *Poz*, paralleling the reduction in *Zfp* (**Figure 5J**). Furthermore, yeast two–hybrid assays further confirmed the interaction between ZFP and either RS27ae or POZ (**Figure 5K**). Together, these results indicate that ZFP could serve as an important transcriptional node potentially regulating male germ cell differentiation in *S. japonicum*, mediated by the Hedgehog signal pathway. Furthermore, the observed effects following *Zfp* RNAi on vitellarium development and egg production in females suggest that *Zfp* could also be involved in male-female interaction of *S. japonicum*, directly or indirectly controlling male-induced processes in the paired female.

### 2.6 Male *Lim^+^* muscle cells influence female development and egg production

Given the sustained expansion of muscle cell populations in adult males and their near absence in paired females (**Figure 6A**), we hypothesized that male muscle cells may provide physical support and other unidentified roles during pairing (**Figure 6B** and Supplementary **Table 7**). Interestingly, we found *Lim,* which is a transcription factor known to regulate muscle motility and cytoskeletal organization in mammals^16^, highly expresses in male muscle cells after pairing. Integration of single–cell and spatial transcriptomics, combined with FISH and WISH analyses, we noted that *Lim*⁺ cells were markedly enriched in the tegument—particularly within the male gynecophornal canal—with lower expression also detected in other somatic tissues (**Figure 6C, 6D and 6F**).

**Figure 6.**
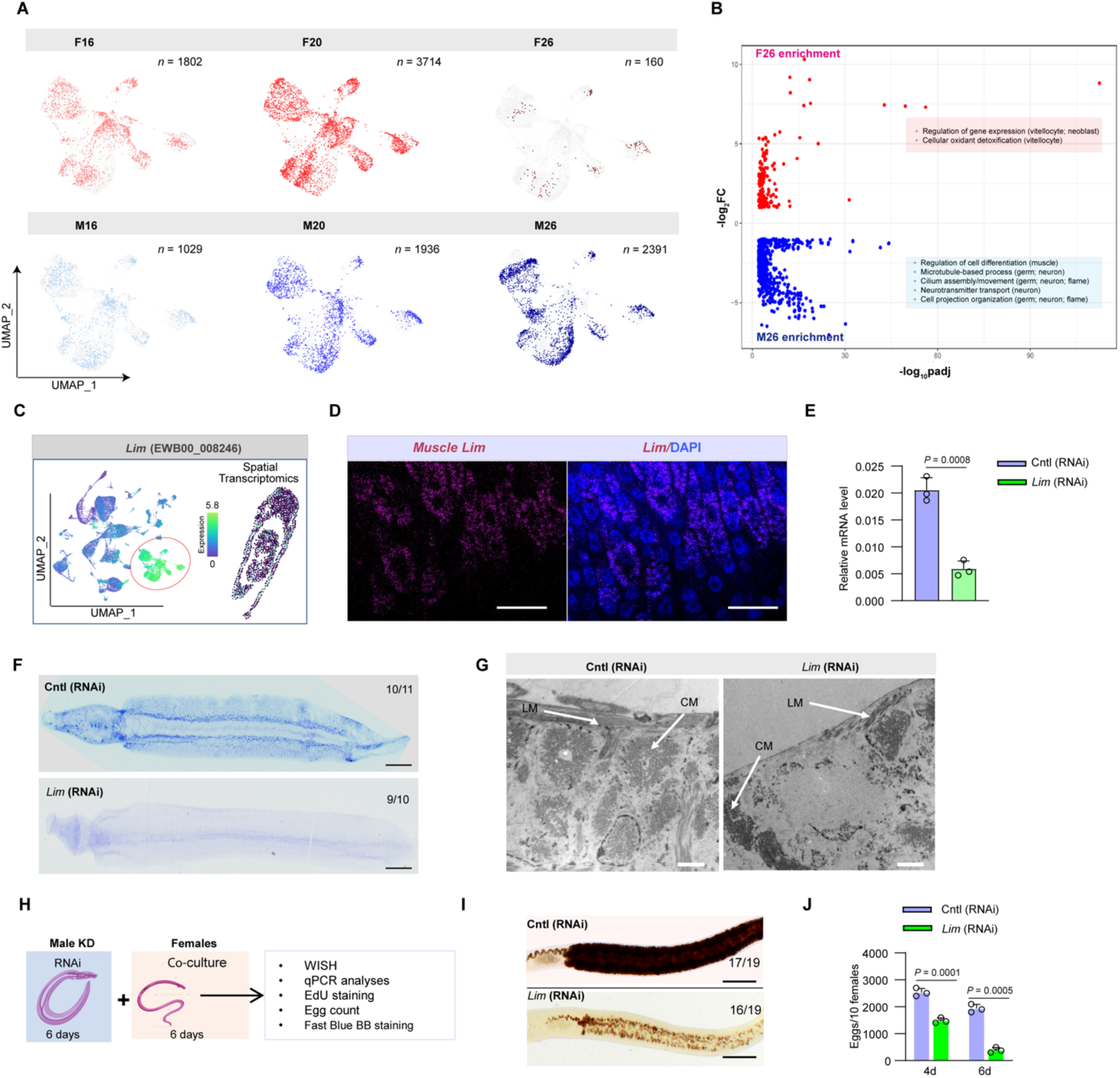
Functional roles of muscle *Lim^+^* cells in male *S. japonicum*. **(A)** UMAP plot of muscle cells across different stages between males and females. (**B**) GO analysis of the biological functions of different genes from different type cells between adult male (M26) and adult female (F26). (**C**) UMAP plot and spatial transcriptomic results indicated *Lim+* cell localization in muscle. **(D)** FISH analysis of *Lim^+^* cell localization in male muscle. Scale bars = 10 μm. **(E)** RT-qPCR analysis of the transcript level *of Lim*^+^ in dsRNA-treated males. Parasites were collected at 6 days of post treatment and then subject to RNA isolation for RT-qPCR analysis. Data illustrate representative results indicating mean and standard deviation obtained from an experiment conducted in triplicate under two biological replicates. (**F**) WISH analysis of *Lim* distributions in control and dsRNA-treated males. A representative image is shown from 9-11 males. Numbers given indicate the fraction of worms that were similar with respect to the phenotype in relation to the total number of worms examined. Scale bars = 200 μm. (**G**) TEM analysis of morphological alterations in *Lim* KD males. LM: longitudinal muscle; CM: Circular smooth muscle. Data are representative results from at least 10 worms investigated in three independent experiments. Scale bars = 2 μm. (**H**) Scheme of functional analysis of *Lim*^+^ cell in adult males and females. (**I**) FastBlue BB staining showing vitellaria in female worms co-cultured with *Lim* KD males. Female schistosomes were stained with Fast Blue BB on days 12 of post co-culture with *Lim* KD males. Data shown is a representative image from 16-19 females. Numbers given indicate the fraction of worms that were similar with respect to the phenotype in relation to the total number of worms examined. Scale bars = 200 μm. (**J**) Egg production significantly decreased in females co-cultured with *Lim* KD males. Data illustrate representative results as mean ± standard error derived from the experiments of triplicates.

Next, we used RNAi to knockdown (KD) *Lim* in males as validated by qPCR and WISH (**Figure 6E and 6F**). Transmission electron microscopy analyses revealed that *Lim* suppression led to reduced muscle integrity in the tegument (**Figure 6G**). To examine how impaired muscle function in males influences female development, we co–cultured *Lim* KD males with untreated females (**Figure 6H**). Eventually, male and female in the control dsRNA treatment showed active twining together under in vitro culture conditions, although they were unable to fully pair at the gynecophoral canal. Expectedly, *Lim* KD males exhibited less physical activities for hugging females. FastBlue BB staining analysis of co-cultured females showed altered patterns of vitellarium (**Figure 6I**), suggesting that vitellarium development could be influenced by male muscle cells. Next, we monitored egg production during dsRNA treatment and found that the number of eggs were significantly reduced in the females co–cultured *Lim* KD males (**Figure 6J**). Overall, these results display that *Lim*⁺ muscle cells in males not only play an important role in maintaining tegumental integrity but also affect vitellarium development and egg production for females.

### 2.7 Germline stem cell lineage across *S. japonicum* development and sexual maturation

Having defined the TFs that regulate male and female reproductive maturation, we next turned to the contribution of stem cells to parasite development and sexual maturation. Using *Ago2,* a conserved marker of pluripotent stem cells^17^, we re-clustered stem cell populations across differentially development stages, identifying 3,582, 2,538, and 2,255 *Ago2⁺* cells at D16, D20, and D26, respectively. These *Ago2⁺* populations exhibited substantial heterogeneity, encompassing somatic stem cells of muscle, neuron, tegument, tegument–progenitor, parenchyma, gut and neoblast lineages. Such somatic stem cells were present at all stages, both in juveniles and adults, suggesting that tissue–autonomous progenitors sustain organ homeostasis throughout the parasite’s life (**Figure 7A** and **7B**). Since germline stem cells (GSCs) form a distinct lineage responsible for gametogenesis^18^, we further analyzed GSC lineage in *S. japonicum* by re-clustering *Ago2⁺*cells that express *Nanos-1*. *Ago2^+^/Nanos-1^+^* cell populations were detectable from juveniles to adults, suggesting that schistosomula as early as D16 already possess GSCs capable of driving sexually dimorphic development (**Figure 7A** and **7B**, Supplementary **Table 8**).

**Figure 7.**
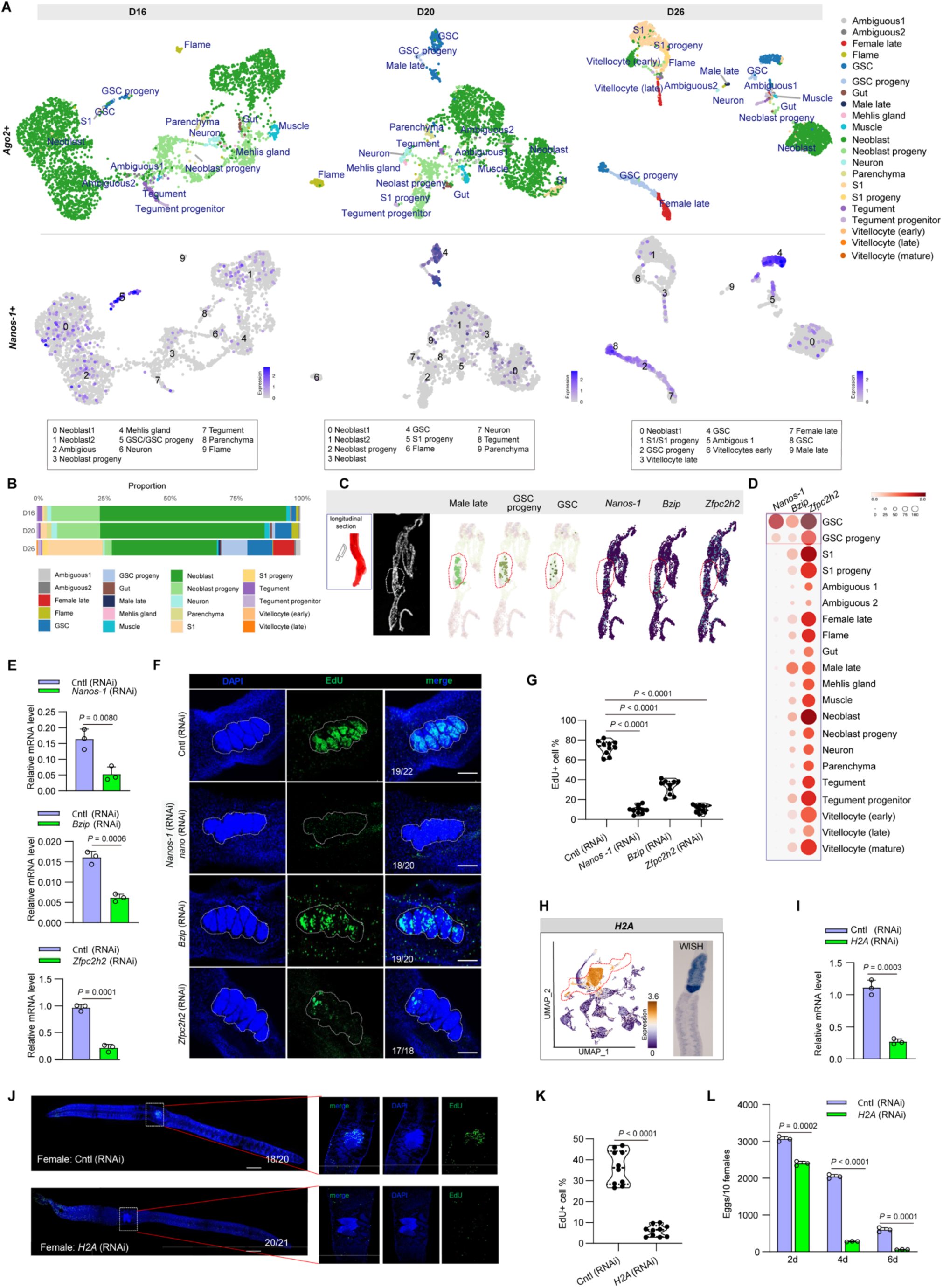
Germline stem cell lineage and functional regulation during *S. japonicum* sexual maturation. (**A**) UMAP plots of *Ago2^+^* subclusters (upper panel), and *Nanos-1* expression in *Ago2+* subclusters (lower panel) across developmental stages. (**B**) The proportions of *Ago2^+^* cell type at different stages (D16, D20, D26). (**C**) Spatial transcriptomic data showed the localization of *Nanos-1*, *Bzip*, and *Zfpc2h2* that co-localize with male late germ cells, GSCs, and GSC progeny. (**D**) scRNA-seq expression profiles of *Nanos-1*, *Bzip* and *Zfpc2h2* in different cell types. (**E**) RNAi inhibition of *Nanos-1* and *Bzip* decreased their transcript levels in adult males. Parasites were collected at 6 days of post treatment and then subject to RNA isolation for RT-qPCR analysis. Data illustrate representative results from two biological replicates. The results indicate mean and standard deviation obtained from an experiment conducted in triplicate. (**F**) Inhibition of *Nanos-1* or *Bzip* influence cell proliferation in testis of male schistosome. The shown image is a representative result from 17-22 males. Numbers given indicate the fraction of worms that were similar with respect to the phenotype in relation to the total number of worms examined. Scale bars = 100 μm. (**G**) Quantification of EdU⁺ cells in testes from *Nanos-1*, *Bzip* or *Zfpc2h2* KD males. *n* = 10. (**H**) UMAP and WISH analyses of localization of *H2a* in adult females. (**I**) Suppression of *H2a* decreased its transcript level in adult females. Parasites were collected at 6 days of post treatment and then subject to RNA isolation for RT-qPCR analysis. Data illustrate representative results indicating mean and standard deviation obtained from an experiment conducted in triplicate from two biological replicates. (**J**) Representative EdU staining analyses for ovarian and vitellarium proliferation in *H2a* KD females. Data shown is a representative image from 18-21 females. Numbers given indicate the fraction of worms that were similar with respect to the phenotype in relation to the total number of worms examined. Scale bars = 300 μm. (**K**) Quantification of EdU⁺ cells in testes from *H2a* KD females. *n* = 10. (**L**) Egg production was significantly reduced in *H2a* KD females. Data illustrate representative results as mean ± standard error derived from the experiments of triplicates.

To uncover the molecular program governing germ-cell differentiation, we focused on TFs with sustained high expression across these stages, including *Nanos-1* (EWB00_002258), *Bzip* (EWB00_005543), and *Zfpc2h2* (EWB00_004260). Single–cell expression profiles revealed significant enrichment of these TFs in GSCs and GSC progeny (**Figure 7D**), and spatial transcriptomics confirmed their localization within GSC niches of males (**Figure 7C**). RNAi–mediated suppression of each TF significantly reduced its transcript levels (**Figure 7E**). Notably, *Nanos-1* and *Zfpc2h2* suppression most severely impaired testicular cell proliferation (**Figure 7F** and **7G**), suggesting their critical roles in maintaining male GSC differentiation.

We also observed consistently high expression of histone *H2A* (EWB00_006186), a core nucleosome component that require for chromatin organization and epigenetic regulation^19^, in *Ago^+^*/*Nanos-1^+^*cell populations throughout worm development, particularly in GSC. ScRNA-seq data, together with WISH, confirmed pronounced enrichment of *H2A* in gonadal tissues including ovary and vitellocytes as well as neoblasts (**Figure 7H**). RNAi-mediated suppression of *H2A* in females (**Figure 7I**) significantly reduced egg production (**Figure 7J**), accompany with decreased cell proliferation in ovary and vitellarium (**Figure 7K** and Supplementary **Figure 10**). These results suggest that H2A is essential for female ovarian development and H2A–containing nucleosomes may maintain genetic and epigenetic fidelity in germ stem cells^20^.

### 2.8 Cellular plasticity between schistosome species

Schistosomiasis is caused by distinct parasite species in different geographical regions. *S. japonicum* and *S. mansoni*, two of the most prevalent species, differ markedly in adult size, egg morphology, and fecundity ^21,22^. To reveal the cellular basis of these species–specific traits, we performed an integrated comparative analysis of our *S. japonicum* single–cell atlas and a published *S. mansoni* dataset.

After aligning the two single–cell datasets (**Figure 8A** and Supplementary **Figure 11A**), we found that both species contain largely homologous cell clusters (Supplementary **Figure 11B**). Cross–species correlation analysis revealed a clear hierarchy of conservation: undifferentiated stem and progenitor cells, such as neoblasts, germline stem cells (GSCs), and S1 cells, were highly conserved (*r* ≥ 0.80; **Figure 8B and 8C** and Supplementary **Figure 11C**), reflecting that they may share developmental programs ^23^; in contrast, terminally differentiated cells such as late-stage male germ cells (r = 0.60), gut, neurons, tegument and parenchyma exhibited more pronounced transcriptional divergence (*r* < 0.75), demonstrating that species-specific adaptations are most prominent at the terminal stages of cellular differentiation.

**Figure 8.**
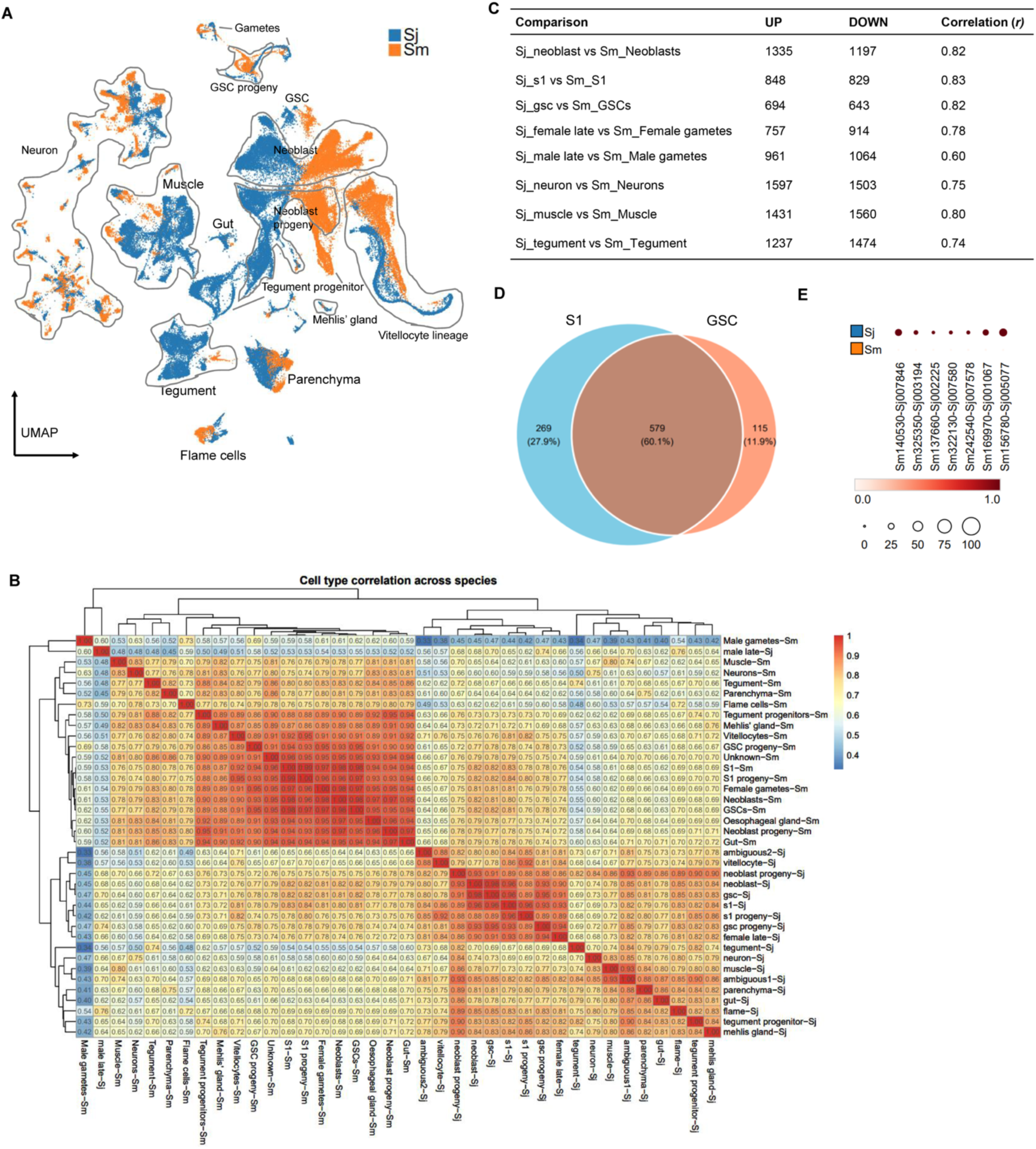
Cross-species analyses of scRNA-seq data between *S. japonicum* and *S. mansoni*. (**A**) Clustering of integrated scRNA-seq data for *S. japonicum* (Sj) and *S. mansoni* (Sm). Major cell types are indicated. (**B**) Pairwise correlation analysis of major cell types between two species, with Pearson’s correlation coefficient (*r*) labelled. (**C**) Numbers of differentially expressed genes (DEGs) in selected cell types between Sj and Sm. (**D**) Venn diagram showing the overlap of upregulated genes between S1 and germline stem cells (GSCs). (**E**) Dot plot indicating the preferential gene expression in *S. japonicum* compared with *S. mansoni*.

To define the molecular basis of these differences, we performed pseudo–bulk differential expression analyses. Consistent with the conservation hierarchy, stem cells contained far fewer differentially expressed genes (DEGs) than specialized cell types (e.g., 1,337 DEGs in GSCs versus 3,100 DEGs in neurons; **Figure 8C**). Focusing on DEGs in GSCs and S1 cells, which are likely involved in the divergent egg–output phenotype (Supplementary **Table 9**), we found that 60% of genes upregulated in *S. japonicum* were shared between these two stem–cell populations (**Figure 8D**). Module score and GO enrichment analysis revealed that these shared genes (579 genes) primarily drive protein ubiquitination and biosynthesis (Supplementary **Table 9**), pointing to a synchronized transcriptional program that enhances biomass production across reproductive lineages. This shared set included key regulators of cell division and biomass production, such as a replication factor (Sm140530-Sj007846), a kinesin-like protein (Sm137660-Sj002225), the chromatin assembly factor 1 (Sm322130-Sj007580), and a ribosome biogenesis protein (Sm156780-Sj005077). These genes were highly and preferentially expressed in *S. japonicum* stem cells but were barely detectable in *S. mansoni* (**Figure 8E, Supplementary Figure 11D and 11E**). The remaining 40% of upregulated genes showed lineage-specific functions, with S1-specific genes enriched in transport in the vitellocyte lineage, and GSC-specific genes associated with muscle contractile components (Supplementary **Figure 11C)**. Collectively, these findings indicated a stem–cell–centric model of evolutionary divergence in *Schistosoma*: transcriptional rewiring of growth and biosynthetic programs confers a hyper–proliferative advantage to *S. japonicum* stem cells, which in turn likely drives its high–fecundity life–history strategy.

## 3 Discussion

The schistosome life cycle encompasses asexual proliferation within intermediate hosts and sexual reproduction in definitive hosts^1^. The transition to sexual maturity represents a pivotal event, linking developmental reprogramming with reproduction and disease transmission. Here, we reported an integrated single-cell and spatial transcriptomics data sets that resolve the cellular architecture and molecular trajectories underlying sexual development in *S. japonicum*. The resulting dataset provides a systematic framework for delineating tissue differentiation and reproductive specialization across key developmental stages. Anatomical mapping via whole–mount in situ hybridization and fluorescence in situ hybridization localized the majority of identified cell types, with spatial transcriptomics offering independent validation. Notably, several cell types such as muscle, neurons, neoblasts, and tegumental cells, displayed significant heterogeneity, which is consistent with previous reports in *S. mansoni*^11^. Together, these comprehensive results established a high–resolution cell atlas for *S. japonicum,* which will enhance understanding of schistosome biology.

Our single–cell atlases of *S. japonicum* across adult stages reveal that sexual maturation is orchestrated through dynamic cellular remodelling, with germline stem cell (GSC) differentiation tightly linked to male–female pairing^6,24^. Pre–paired worms (16 days post–infection) of both sexes share similar somatic–cell compositions, whereas the emergence of female–specific vitellocytes and the expansion of germline–associated populations (S1 cells, GSCs and their progeny) coincide with pairing (20 dpi). By full sexual maturity (26 dpi), males and females follow markedly divergent cellular trajectories. Collectively, this work establishes—for the first time—a dynamic cellular map across the critical phases of sexual maturation and egg production. These findings provide a direct cellular framework for elucidating the molecular and metabolic program that underlie sex–specific development and male–female interaction in schistosomes.

Male–female pairing has long been recognized as central to female sexual maturation and egg production in schistosomes. A previous study in *S. mansoni* reported that even amputated male segments lacking reproductive organs can induce localized development of female vitellaria or ovaries via pairing^25^. Consequently, identifying key molecules that drive sexual development has been a major focus in the field. Several proteins such as transforming growth factor β receptor ^26^, mitogen-activated protein kinase (MAPK) ^27^ pathways, Venus kinase receptors (SmVKRs)^28^, syk tyrosine kinase 4 (SmTK4)^29^, certain phosphoproteins ^30^, and allatostatin A receptor-like protein^31^ have been implicated in schistosome male–female interaction and sexual maturation. More recently, β–alanyl–tryptamine, a male–derived pheromone, was identified as a key mediator of female maturation in *S. mansoni*^32^. Nevertheless, the molecular regulation through which pairing induces female maturation still remains poorly defined, especially in *S. japonicum*.

In the present study, leveraging single–cell transcriptomics, we found that distinct male cell populations such as *Nt3gfr^+^* parenchyma/testis cells, *Zfp*^+^ and *Fbp3*^+^ testis cells and *Lim*^+^ muscle cells can regulate cell proliferation in target tissues. To determine whether these cell populations in males could influence sexual maturation and egg production in females, we co-cultured dsRNA treated males with untreated females. We chose AB169 (1640) as culture medium that had been shown to support worm activity and egg production in vitro ^33^. Notwithstanding, current in vitro culture conditions and medium cannot fully replicate the physical environment and/or provide the essential nutrition present in vivo that support stable and complete male-female pairing. We must acknowledge that males and females did not pair stably or completely under our co-culture conditions; nevertheless, male and female showed twine and partially pair in controls while these phenomena were to some extend weakened in females co-cultured with dsRNA treated males. Consequently, we focused primarily on the effect of co–culture on vitellocyte development and egg production in females. As compared to the control RNAi treatment, in which males were co-cultured with females, we found that knockdown of *Nt3gfr^+^* parenchyma/testis cells, *Zfp*^+^ and *Fbp3*^+^ testis cells and *Lim*^+^ muscle cells resulted in the degenerated vitellarium in co-cultured females. This suggests that hormones, sperm and/or other unidentified testis–derived factors may contribute to vitellarium development and egg production. Given the dynamic cellular remodelling and complex gene–expression changes during sexual development, we deduce that male-induced female sexual maturation through pairing could be a multifaceted mechanism including multiple factors and multiple processes at multiple development stages that especially contribute to female sexual maturation ^34,35^ This is in line with previous findings in *S. mansoni* ^36^ ^37^.

Unlike free-living planarians such as *Schmidtea mediterranea*, schistosomes have lost regenerative capacity despite their shared flatworm ancestry. Our analysis of stem–cell populations across developmental stages reveals a heterogeneous compartment comprising both pluripotent and lineage–primed progenitors—including germline, muscle, neural and Mehlis’ gland progenitors—each following distinct developmental trajectories. This diversity suggests that *Schistosoma* may maintain a reservoir of stem cells that support multiple tissue types, a feature that may enhance its adaptability to host environments and its ability to evade immune responses and ensure transmission ^38^ Therefore, targeting key regulators of stem–cell differentiation could represent an alternative strategy against this disease. Moreover, cross–species integration and comprehensive comparative analyses highlighted species–specific transcriptional programs within the germline and vitellocyte stem cell lineages. We recognize that technical heterogeneities such as the use of different library preparation kits and varying versions of the Cell Ranger pipeline are inherent challenges comparing omics data sets. Our findings appear to reflect genuine biological divergence rather than technical artifacts. First, we implemented a robust homology-based integration using the Harmony algorithm ^39^, which is specifically designed to harmonize datasets across different platforms while preserving biological variation. Second, the successful alignment of major cell lineages and stable expression profiles of fundamental markers (e.g., nanos and vasa) across both species demonstrate a high biological signal-to-noise ratio. Given that *S. japonicum* and *S. mansoni* diverged from a common ancestor millions of years ago ^40^, the observed transcriptional shifts likely represent this evolutionary divergence between these species that far outweighs the subtle biases introduced by the applied experimental protocols.

These divergent molecular signatures likely underpin the distinct life–history traits—such as higher fecundity, unique egg morphology and larger adult size—observed in *S. japonicum*. Notably, the *S. japonicum*-specific upregulated program in GSCs was significantly enriched in myofibril and contractile muscle fiber components (e.g., tropomodulin and paramyosin). In schistosomes, the reproductive tract is structurally integrated with specialized musculature required for egg transport and the maintenance of prolonged pairing. The simultaneous reinforcement of the germline “engine” and the muscular “pump” suggests a coordinated evolutionary adaptation: *S. japonicum* has not only accelerated its stem cell proliferation but also upgraded the mechanical infrastructure necessary to support its massive daily egg output. Consequently, these findings may illuminate how molecular evolution within the reproductive stem–cell compartment and its supporting anatomical niche facilitates ecological adaptation and niche specialization in different schistosome species. In summary, our studies established a comprehensive framework for understanding stem–cell biology and differentiation in schistosomes, and it provides fundamental insights into germline development and reproductive maturation.

## Data and code availability

The raw sequence data of spatial transcriptomic profiling have been deposited in the Genome Sequence Archive in National Genomics Data Center of China National Center for Bioinformation / Beijing Institute of Genomics, Chinese Academy of Sciences (GSA: CRA041784) that are publicly accessible at https://ngdc.cncb.ac.cn/gsa. The scRNA-seq raw data have been deposited in the NCBI Sequence Read Archive database under Bioproject accession: PRJNA1417122 of Sequence Read Archive (SRA) with accession numbers: SRR37125568, SRR37125569, SRR37135933, SRR37135934, SRR37125570, and SRR37125571 [https://www.ncbi.nlm.nih.gov/sra/?term=PRJNA1417122]. The comprehensive datasets are freely available as an online database (https://schisto.xyz/sj-atlas/). All custom code and computational pipelines used to perform the analyses described in this study are publicly available in the following GitHub repository: https://github.com/zglu/Sj-Atlas-Code.

## Acknowledgments

This study was, in part or in whole, supported by National Natural Science Foundation of China (Grants No. 32361133554 and 32573460 to GC) and the Fundamental Research Funds for the Central Universities (PA2024000471, PA2023000254, PA2022000010, PA2021000155, and PA2020000431 to GC). The study is also sponsored by Shanghai Tongji University Education Development Foundation.

## Author contributions

Conceptualization, G.C. and Z.L.; methodology, X.W., Z.L., S.L., X.L., Y. Z., L.L., and P.D.; software, Z.L.; formal analysis, Z.L., X.W., Y.Z., L.L., and P.D.; resources, Y.Z., and G.C.; investigation, X.W., Z.L., S.L., X.L., Y.Z., and L.L.; visualization, X.W., and Z.L.; technical assistance, Z.L. and P.D.; project administration, G.C.; supervision, G.C.; writing—original draft, G.C. and Z.L; writing—review & editing, G.C., Z.L. and O.Z.

## Declaration of interests

The authors declare no competing interests.

## **4** Materials and methods

### Animal and parasites

Male BALB/c mice (6–8 weeks of age) and male New Zealand rabbits (3 months of age) were obtained from the Shanghai Experimental Animal Center, Chinese Academy of Sciences. *Schistosoma japonicum* cercariae were freshly harvested from infected snails and provided by the National Institute of Parasitic Diseases at the Chinese Center for Disease Control and Prevention. Percutaneous infection was carried out by exposing the shaved abdominal skin of each animal to the cercariae, with target doses of 200 ± 10 cercariae per mouse and 2,000 ± 50 cercariae per rabbit. The animal protocols were approved by the Institutional Animal Care and Use Committee of Tongji University School of Medicine (No. TJAA00824502)

### Parasite collection and culture

*S. japonicum* worms were isolated from the hepatic portal system of infected mice by perfusion with citrate-supplemented PBS pre-equilibrated to 37°C. After perfusion, parasites were repeatedly rinsed with RPMI 1640 (Gibco) supplemented with 1% penicillin-streptomycin and transferred to ABC169 medium for maintenance at 37°C with 5% CO₂. The medium was changed every 1–3 days.

### Single-cell RNA sequencing (scRNA-seq)

*S. japonicum* worms at 16 dpi, 20 dpi and 26 dpi were dissociated into single-cell suspensions and loaded by a SONY SH800S cell sorter as described previously ^41^. Dead cells were removed using the Dead Cell Removal Kit (Miltenyi Biotec). Single-cell RNA sequencing was performed using 10 × Genomics Chromium platform, in accordance with our previously published procedure ^41^.

### Mapping of scRNA-seq reads

Sequencing reads were aligned to a custom-built *S. japonicum* reference genome derived from the assembly available under BioProject PRJNA520774 (WormBase ParaSite), to which mitochondrial gene sequences (NCBI RefSeq NC_002544.1, designated “Sj”) were manually appended. The alignment was performed using Cell Ranger (version 5.0).

### Reference gene information

Orthologs between *S. mansoni* (v7) and *S. japonicum* (HuSjv2), along with their protein sequences and GO annotations, were extracted from WormBase ParaSite (https://parasite.wormbase.org/) ^42^ using the BioMart tool (access date: 2024-05-11). Single-cell marker genes were validated against published datasets from *S. mansoni* adults (filtered at p_val < 1×10⁻²⁰)^11^ and schistosomula ^9^ (filtered by myAUC > 0.7). *S. japonicum* genes were assigned using the ortholog relationships described above.

### Identification of *S. japonicum* transcription factors

First, protein sequences for the HuSjv2 assembly were downloaded from WormBase Prasite version 16 and subjected to KEGG ortholog mapping using the KEGG Automatic Annotation Server (KAAS) with the Bi-Directional Best Hit (BBH) method (https://www.genome.jp/kegg/kaas/; access date: 2021-11-27). This mapping resulted in a list of orthologous genes that were further classified as eukaryotic transcription factors (TFs) based on the KEGG/TRANSFAC 7.0 database (https://www.kegg.jp/kegg-bin/get_htext?ko03000.keg). Second, the sequences of identified TFs were subjected to protein domain analysis using InterProScan v5.77-108.0 against all default databases including Pfam, SMART, and CDD (https://www.ebi.ac.uk/interpro/). Only proteins harboring recognized DNA-binding domains (e.g., zinc finger, homeobox, bHLH, bZIP, Forkhead, and HMG box) were retained as *S. japonicum* TFs.

### ScRNA-seq data processing and annotation

The filtered matrix files of Cell Ranger were processed using Seurat (v4.3.0; https://satijalab.org/seurat/) by reading gene IDs and filtered by nFeature_RNA 500-5,000 & nCount_RNA 1,000-30,000 (Female26 50000) & percent.mt < 3. Potential doublets were identified and removed using scDblFinder (v1.13.7), ensuring that only singlets were used to subsequent analyses. After normalization using SCTransform (vars.to.regress=”percent.mt”) and PCA dimensionality reduction, all six datasets were integrated using the RPCA method according to Seurat tutorial (https://satijalab.org/seurat/archive/v4.3/integration_rpca). Briefly, the command SelectIntegrationFeatures was run to select top 3,000 variable features and then PrepSCTIntegration, FindIntegrationAnchors with parameters normalization.method = “SCT” and reduction = “rpca”. Data were then integrated with IntegrateData and PCA was run again. A neighborhood analysis was performed using the first 50 principal components, and cell clusters were identified with the default clustering algorithm at a resolution of 2. Cell types for scRNA-seq data were annotated as described in our previous publication ^41^.

### Correlation analyses of global expression

To assess the global expression among all six samples, a pseudo-bulk expression correlation analysis was performed. In each sample, the average gene expression vectors were calculated across all of shared genes using the Seurat AverageExpression function based on normalized counts (the ‘data’ slot). Pearson correlation coefficient was computed for all pairwise sample combinations.

The resulting 6 × 6 correlation matrix was then visualized as a heatmap using the pheatmap package (v1.0.13). Hierarchical clustering was applied to both the rows and columns of the matrix using the ward. D2 method to group samples exhibiting the most similar global expression profiles.

### Pseudo-bulk differential gene expression analysis

To identify genes exhibiting differential expression across developmental stages and/or sexes, we employed a pseudo-bulk differential gene expression (DGE) analysis on the scRNA-seq data. Briefly, raw UMI counts from clustered cells were aggregated to create pseudo-bulk samples using the Seurat Aggregate Expression function, with the parameter group.by = c(“orig.ident”, “main_type”). Differential expression analysis was performed with DESeq2^43^ (v1.42.1) using the aggregated count matrix as input. DGE comparisons were performed using two main strategies: 1) a pooled stage-level comparison (e.g., D20 vs. D16 with sexes combined); and 2) adjacent, sex- and stage-specific comparisons (e.g. F20 vs. F16 and F26 vs. F20, etc.). Significant DEGs were identified at padj < 0.01 and |log₂FC| ≥ 1. All resulting DEGs from these comparisons were compiled into a master list representing genes with dynamic expressions. To investigate the temporal patterns of expression changes within this list, we performed fuzzy clustering using the Mfuzz v2.62.0 package^43^. Briefly, the expression data for these DEGs were extracted from the Seurat object and ordered according to their respective developmental stages (F16, F20, F26, M16, M20, and M26). They were then clustered using the Mfuzz function. The fuzzification parameter *m* was estimated using mestimate, and the number of clusters *c* was set to 8. Finally, the signature score for the genes within each resulting cluster was calculated across the single-cell expression data using the Seurat AddModuleScore function.

### Functional enrichment analysis

GO enrichment analysis was performed on fuzzy-clustered genes using the g:GOSt module of g:Profiler (v e113_eg59_p19_f6a03c19). Terms (adjusted *P* < 0.05 and ≥ 5 annotated genes) were considered significant, and The results of Biological Process were obtained for visualization.

### Protein-protein interaction (PPI) network analysis

STRING v12.0 was used to build a *S. japonicum* PPI network from the input gene set. Interactions with a combined score ≥ 0.400 (medium confidence) were retained based on experimental, co-expression, database, and text-mining evidence. The network was clustered into functional modules and visualized.

### Cross-species comparison of scRNA-seq data between *S. japonicum* and *S. mansoni*

To facilitate robust comparative analysis between the single-cell RNA sequencing datasets of *S. japonicum* and *S. mansoni*, a homology-based integration strategy was implemented. The *S. japonicum* dataset, including day 16 (D16), day 20 (D20), and day 26 (D26), was integrated with published *S. mansoni* data, which included immature females and adult worms of both sexes. Expression matrices were filtered to include only 1:1 orthologous genes (approximately 6,800 genes per species) prior to merging. The combined dataset underwent standard Seurat (v5.0.1) pre-processing, including normalization, variable feature selection, and scaling. Principal Component Analysis (PCA) was performed, retaining 80 principal components (PCs). To account for potential biases introduced by different developmental stages and library preparation protocols, batch correction and data alignment across the two species were achieved using the Seurat IntegrateLayers function with the HarmonyIntegration method (orig.reduction = “pca”, new.reduction = “harmony”). Following integration, cells were clustered using FindClusters (resolution = 0.5), and layers were combined using JoinLayers for differential expression analysis. The integrated object’s metadata retained both the original cluster annotation and a unified main cell type annotation (eg. neoblast for *S. japonicum* and Neoblasts for *S. mansoni*).

To identify genes with differential expression between the two species within specific cell populations (e.g., neoblasts), cell type-specific pseudo-bulk samples were generated. Raw UMI counts corresponding to specific cell groups were aggregated using the Seurat AggregateExpression function (grouped by: group.by = c(“species”, “orig.anno”)) resulting in a total of 130 pseudo-bulk samples based on original cluster annotations. Pparallel aggregation was performed using a unified main cell type annotation for expression correlation analysis. Differential expression analysis of these aggregated raw counts was conducted using the DESeq2 package. Significant DEGs were identified using thresholds of padj < 0.01 and log₂ fold change ≥ 1. Module scores were calculated using AddModuleScore function in Seurat.

### Quality control of OCT-embedded samples

For the OCT-embedded sample, 100–200 μm thick sections were cut and used to extract total RNA using the RNeasy Mini Kit (Qiagen, USA). The RNA integrity number (RIN) was checked by a 2100 Bioanalyzer (Agilent) and samples with RIN ≥ 7 were selected for downstream experiments. Cryosections with a thickness of 10 μm were obtained for hematoxylin and eosin (H&E) (BGI, China) to examine the tissue morphology. Samples were selected for the Stereo-seq experiment.

### Stereo-seq library preparation and sequencing

Adult parasites were collected from infected mice. OCT-embedded samples were prepared and quality control was performed as described previously ^44^. Spatial transcriptomics experiment were performed using the STOmics FF V1.3 gene expression assay kit (BGI, China) according to the manufacturer’s protocol.

### Stereo-seq data processing and cell type annotation

Sequencing data from Stereo-seq spatial transcriptomics were processed using Stereo-seq Analysis Workflow (SAW) v8.0 (https://en.stomics.tech). Briefly, reads corresponding to the chip spatial barcodes were mapped to the HuSjv2 reference genome and annotated to generate a bin1 expression matrix, which was then aligned with the ssDNA-stained microscope image for tissue and cell segmentation. The cell-level expression data were further corrected for cell border expansion using a predefined algorithm. Unsupervised clustering was performed in the analysis. For cell type prediction, we used RCTD v2.2.0 ^45^ using our annotated scRNA-seq data as a reference. The doublet mode was choose, and the cell types were assigned with the highest weight. In order to validate the annotation results, we manually examined the expression of cell-type-specific marker genes in both single-cell and spatial transcriptomic datasets.

### Whole mount of *in situ* hybridization (WISH) and fluorescence *in situ* hybridization

WISH analyses were performed as described previously^41,46–48^. Worms were hybridized overnight at 54–56°C with 100 ng/mL DIG-labeled or control riboprobes. Following washes, samples were blocked (10% horse serum, 2 h, RT) and incubated with anti-DIG-AP Fab fragments (1:2,000; Roche) overnight at 4°C. After TNT washing (2 h, changes every 20 min), worms were equilibrated in AP buffer (100 mM Tris-HCl pH 9.5, 100 mM NaCl, 50 mM MgCl₂, 0.1% Tween-20, 10% polyvinyl alcohol, 10 min) and stained with NBT/BCIP for 3 h to overnight. Stained worms were washed in PBST, post-fixed, and mounted in 80% glycerol for imaging.

For probe preparation, cDNA fragments were amplified from 28-day adult worm total RNA-derived cDNA using gene-specific primers (Supplementary Table 10). Purified PCR products were then used as templates to generate DIG-labeled riboprobes via the MEGAscript Kit (Thermo Fisher Scientific) with DIG-11-UTP (Roche).

For FISH analysis, target gene fragments were amplified using PCR. Following PCR product purification, in vitro transcription was conducted using the MEGAscript kit (ThermoFisher Scientific) according to the manufacturer’s instructions. RNA purification was performed using the protocol described for WISH probe preparation. Probe concentration was quantified via a NanoDrop spectrophotometer (ThermoFisher Scientific). FISH experiments were preformed as described in our previous publication ^44^.

### Schistosome culture, RNAi experiment and RT-qPCR analyses

Worms recovered from mice at 22–24 dpi were sex-separated and cultured in AB169 (1640) medium (37°C, 5% CO₂) ^33^. dsRNA (30 µg/mL) generated using the MEGAscript Kit was added to the medium. For egg-laying assays, dsRNA-treated males (6 days) were washed and co-cultured with untreated females, eggs were counted using inverted microscopy. Parasites were sampled at indicated times for RT-qPCR using ChamQ Universal SYBR qPCR Master Mix on a StepOne Plus system. Primers used are listed in Supplementary **Table 10**. *S. japonicum NADH* (forward primer: 5′-CGAGGACCTAACAGCAGAGG-3′; reverse primer: 5′-TCCGAACGAACTTTGAATCC-3′) was used for normalization, and relative expression was calculated by the 2^-ΔCt^ method ^49^.

### FastBlue BB and EdU staining

Female worms were fixed (4% formaldehyde in PBSTx, 4 h), stained with freshly filtered 1% Fast Blue BB in PBSTx (5 min), washed thrice with PBSTx, cleared in 80% glycerol, and mounted for microscopic imaging. EdU staining was performed as previously described ^50^.

### Yeast two hybrid analysis

The cDNA sequences of ZFP (EWB00_005374), RS27ae (Putative small subunit ribosomal protein S27Ae, EWB00_000191) and POZ (Speckle-type POZ protein-like B, EWB00_003497) were synthesized by Shanghai Saiheng Biotechnology Company and then cloned into the pGBKT7 (ZFP) and pGADT (RS27ae and POZ) vectors, respectively. All positive clones were verified by sequencing, and the corresponding plasmids (pGBKT7-ZFP + pGADT-RS27ae, pGBKT7-ZFP + pGADT-POZ, pGBKT7 + pGBKT7-ZFP, pGBKT7 + pGADT-RS27ae, pGBKT7 + pGADT-POZ, pGADT7-largeT + pGBKT7-p53 (positive control), pGADT7-largeT + pGBKT7-laminC (negative control)) were co-transformed into AH109 yeast cells. Then, the yeast cells were cultured on SD-TL and SD-TLHA plates to evaluate the potential self-activation of each pair. All of combinations showed no self-activation. Next, a single colony containing each pair of combinations was selected and further cultured in SD-TL, SD-TLHA and SD-TLHA+X-α-gal plates. The plates were incubated at 30 °C to evaluate the protein-protein interaction^51^.

### Transmission electron microscopy

Worms were collected, and transmission electron microscopy were performed as previous publication^52^.

## Statistical analysis

All statistical analyses were performed in GraphPad Prism 8. Differences between two groups were assessed using Student’s t-test, whereas comparisons involving multiple groups were evaluated using one-way ANOVA. Statistical significance was defined as *P* ≤ 0.05.

## Supplementary information

**Figure S1.** Identification of cell populations in *S. japonicum*. (**A**) Marker genes in different cell types. (**B**) UMAP showing different cell types. (**C**) Validation of anatomical localizations of representatively identified tissue cells. Red dashed lines on UMAPs highlight the relevant cell cluster. Data are representative results of 20-30 investigated worms. Scale bars = 200 µm.

**Figure S2.** Spatial transcriptomic analyses of *S. japonicum*. (**A**) Spatial transcriptomic marker genes in different cell types. (**B**) Spatial transcriptomic analyses of representative cell types within paired adult parasites.

**Figure S3.** The heterogeneity of neuron cells. Heatmap of neuronal maker genes (**A**) and A dot-plot summarizing the genes highly expressing in each cluster (**B**). (**C**) Double FISH analysis indicated the combination of neuronal cluster-specific markers for neuronal cell population. EWB00_008401, Synaptic vesicle membrane protein VAT-1 isoform 1; EWB00_008189, Calcium uptake protein 2 isoform 2; EWB00_004539, Tyrosine-protein kinase.

**Figure S4.** The heterogeneity of muscle cells. (**A**) Heatmap of muscle maker genes and (**B**) A dot-plot summarizing the genes highly expressing in each cluster.

**Figure S5.** The heterogeneity of tegument cells. (**A**) Heatmap of tegument maker genes and (**B**) A dot-plot summarizing the genes highly expressing in each cluster.

**Figure S6** The heterogeneity of parenchyma cells. (**A**) Heatmap of parenchyma maker genes and (**B**) A dot-plot summarizing the genes highly expressing in each cluster.

**Figure S7** Transcriptomic dynamics during *S. japonicum* sexual maturation. (**A**) Gene expression correlations between different samples. (**B**) UMAP showing the signature score from clustered genes in different cell types between males and females. M, male; F, female.

**Figure S8** EdU staining analysis of cell proliferation in the vitellocytes of *Zeb2* KD females. Data are shown a representative image from 8-10 females. Numbers given indicate the fraction of worms that were similar with respect to the phenotype in relation to the total number of worms examined. Scale bars = 60 μm.

**Figure S9** Ovarian proliferation remained unaffected in *Zfp* and *Fbp3* KD females. Data are shown a representative image from 17-21 females. Numbers given indicate the fraction of worms that were similar with respect to the phenotype in relation to the total number of worms examined. Scale bars = 100 μm.

**Figure S10** EdU staining analysis of cell proliferation in the vitellocytes of *H2a* KD females. Data are shown a representative image from 18-21 females. Numbers given indicate the fraction of worms that were similar with respect to the phenotype in relation to the total number of worms examined. Scale bars = 50 μm.

**Figure S11** Comparison of scRNA-seq data between *S. japonicum* and *S. mansoni.* (**A**) Clustering of integrated scRNA-seq data for *S. japonicum* and *S. mansoni*. (**B**) Unsupervised clustering and source of origin in each cluster. (**C**) Cross-species correlation analysis for identified cell types. (**D**) Dot plot showing cell type expression of identified key genes. The three modules represent shared upregulated genes in both GSC and S1 (Shared_Up, 60%), and lineage-specific upregulated genes (GSC-only and S1-only, 40%) corresponding to Fig 8D and Supplementary Table 9. (**E**) Screenshots showing the query results from the schistosome expression database.

**Supplementary Tables**

**Table S1** The list of cluster maker genes in each cell population.

**Table S2** The list of marker genes for different cell types from spatial transcriptomics.

**Table S3** The list of marker genes for Figure S2B.

**Table S4** The list of identified genes in different clusters by fuzzy clustering analysis and their GO enrichment results.

**Table S5** The list of differently expressed genes between D20 and D16.

**Table S6** The list of identified transcription factors in *S. japonicum*.

**Table S7** The list of differentially expressed genes between F26 and M26.

**Table S8** The list of maker genes of *Ago2^+^* cell populations in D16/D20/D26.

**Table S9** The lists of differentially expressed genes of S1 and germinal stem cells between *S. mansoni* and *S. japonicum*.

**Table S10** The list of primers used in the present study.

## Notes

### Competing Interest Statement

The authors have declared no competing interest.

### Summary of Updates

Discussion had been changed. Fig1 revised. Fig 8 revised.

## References

1 Buonfrate, D., Ferrari, T. C. A., Adegnika, A. A., Russell Stothard, J. & Gobbi, F. G. Human schistosomiasis. Lancet 405, no.10479 (2025): 658–670, 10.1016/S0140-6736(24)02814-9

2 Cotton, J. A. & Doyle, S. R. A genetic TRP down the channel to praziquantel resistance. Trends Parasitol 38, no.5 (2022): 351–352, 10.1016/j.pt.2022.02.006

3 Cheng, G. F. et al. Proteomic analysis of differentially expressed proteins between the male and female worm of *Schistosoma japonicum* after pairing. Proteomics 5, no. 2 (2005): 511–521, 10.1002/pmic.200400953

4 LoVerde, P. T. & Chen, L. Schistosome female reproductive development. Parasitol Today 7, no. 11 (1991): 303–308, 10.1016/0169-4758(91)90263-n

5 Grevelding, C. G., Sommer, G. & Kunz, W. Female-specific gene expression in *Schistosoma mansoni* is regulated by pairing. Parasitology 115, no. 6 (1997): 635–640, 10.1017/s0031182097001728

6 Kunz, W. Schistosome male-female interaction: induction of germ-cell differentiation. Trends Parasitol 17, no. 5 (2001): 227–231, 10.1016/s1471-4922(01)01893-1

7 Attenborough, T. et al. A single-cell atlas of the miracidium larva of *Schistosoma mansoni* reveals cell types, developmental pathways, and tissue architecture. Elife 13 (2024): RP95628, 10.7554/eLife.95628

8 Diaz Soria, C. L., et al. Single-cell transcriptomics of the human parasite *Schistosoma mansoni* first intra-molluscan stage reveals tentative tegumental and stem-cell regulators. Sci Rep 14, no. 1 (2024): 5974, 10.1038/s41598-024-55790-3

9 Diaz Soria, C. L., et al. Single-cell atlas of the first intra-mammalian developmental stage of the human parasite *Schistosoma mansoni*. Nat Commun 11, no. 1 (2020): 6411, 10.1038/s41467-020-20092-5

10 Li, P. et al. Single-cell analysis of *Schistosoma mansoni* identifies a conserved genetic program controlling germline stem cell fate. Nat Commun 12, no. 1 (2021): 485, 10.1038/s41467-020-20794-w

11 Wendt, G. et al. A single-cell RNA-seq atlas of *Schistosoma mansoni* identifies a key regulator of blood feeding. Science 369, no. 6511 (2020): 1644–1649, 10.1126/science.abb7709

12 Morales-Vicente, D. A. et al. Single-cell RNA-seq analyses show that long non-coding RNAs are conspicuously expressed in *Schistosoma mansoni* gamete and tegument progenitor cell populations. Front Genet 13 (2022): 924877, 10.3389/fgene.2022.924877

13 Moescheid, M. F. et al. The retinoic acid family-like nuclear receptor SmRAR identified by single-cell transcriptomics of ovarian cells controls oocyte differentiation in *Schistosoma mansoni*. Nucleic Acids Res 53, no. 4 (2025): gkae1228, 10.1093/nar/gkae1228

14 El-Shehabi, F., Taman, A., Moali, L. S., El-Sakkary, N. & Ribeiro, P. A novel G protein-coupled receptor of *Schistosoma mansoni* (SmGPR-3) is activated by dopamine and is widely expressed in the nervous system. PLoS Negl Trop Dis 6, no. 2 (2012): e1523, 10.1371/journal.pntd.0001523

15 Buddenborg, S. K., Lu, Z., Sankaranarayan, G., Doyle, S. R. & Berriman, M. The stage- and sex-specific transcriptome of the human parasite *Schistosoma mansoni*. Sci Data 10, no. 1 (2023): 775, 10.1038/s41597-023-02674-2

16 Kadrmas, J. L. & Beckerle, M. C. The LIM domain: from the cytoskeleton to the nucleus. Nat Rev Mol Cell Biol 5, no. 11 (2004): 920–931, 10.1038/nrm1499

17 Ngondo, R. P. et al. Argonaute 2 is required for extra-embryonic endoderm differentiation of mouse embryonic stem cells. Stem Cell Reports 10, no. 2 (2018): 461–476, 10.1016/j.stemcr.2017.12.023

18 Lehmann, R. Germline stem cells: origin and destiny. Cell Stem Cell 10, no. 6 (2012): 729–739, 10.1016/j.stem.2012.05.016

19 Kornberg, R. D. Chromatin structure: a repeating unit of histones and DNA. Science 184, no. 4139 (1974): 868–871, 10.1126/science.184.4139.868

20 Kornberg, R. D. & Lorch, Y. Primary role of the nucleosome. Mol Cell 79, no. 3 (2020): 371–375, 10.1016/j.molcel.2020.07.020

21 Rheinberg, C. E. et al. *Schistosoma haematobium*, *S. intercalatum*, *S. japonicum*, *S. mansoni*, and *S. rodhaini* in mice: relationship between patterns of lung migration by schistosomula and perfusion recovery of adult worms. Parasitol Res 84, no. 4 (1998): 338–342, 10.1007/s004360050407

22 Gui, M., Kusel, J. R., Shi, Y. E. & Ruppel, A. *Schistosoma japonicum* and *S. mansoni*: comparison of larval migration patterns in mice. J Helminthol 69, no. 1 (1995): 19–25, 10.1017/s0022149x0001378x

23 Tarashansky, A. J. et al. Mapping single-cell atlases throughout Metazoa unravels cell type evolution. Elife 10 (2021): e66747, 10.7554/eLife.66747

24 Erasmus, D. A. A comparative study of the reproductive system of mature, immature and “unisexual” female *Schistosoma mansoni*. Parasitology 67, no. 2 (1973):165–183, 10.1017/s0031182000046394

25 Popiel, I. & Basch, P. F. Reproductive development of female *Schistosoma mansoni* (Digenea: Schistosomatidae) following bisexual pairing of worms and worm segments. J Exp Zool 232, no. 1 (1984): 141–150, 10.1002/jez.1402320117

26 Osman, A., Niles, E. G., Verjovski-Almeida, S. & LoVerde, P. T. *Schistosoma mansoni* TGF-beta receptor II: role in host ligand-induced regulation of a schistosome target gene. PLoS Pathog 2, no. 6 (2006): e54, 10.1371/journal.ppat.0020054

27 Andrade, L. F. et al. Regulation of *Schistosoma mansoni* development and reproduction by the mitogen-activated protein kinase signaling pathway. PLoS Negl Trop Dis 8, no. 6 (2014): e2949, 10.1371/journal.pntd.0002949

28 Vanderstraete, M. et al. Venus kinase receptors control reproduction in the platyhelminth parasite *Schistosoma mansoni*. PLoS Pathog 10, no. 5 (2014): e1004138, 10.1371/journal.ppat.1004138

29 Beckmann, S., Buro, C., Dissous, C., Hirzmann, J. & Grevelding, C. G. The Syk kinase SmTK4 of *Schistosoma mansoni* is involved in the regulation of spermatogenesis and oogenesis. PLoS Pathog 6, no. 2 (2010): e1000769, 10.1371/journal.ppat.1000769

30 Hirst, N. L., Nebel, J. C., Lawton, S. P. & Walker, A. J. Deep phosphoproteome analysis of *Schistosoma mansoni* leads development of a kinomic array that highlights sex-biased differences in adult worm protein phosphorylation. PLoS Negl Trop Dis 14, no. 3 (2020): e0008115, 10.1371/journal.pntd.0008115

31 Wang, J. et al. Dynamic transcriptomes identify biogenic amines and insect-like hormonal regulation for mediating reproduction in *Schistosoma japonicum*. Nat Commun 8 (2017): 14693, 10.1038/ncomms14693

32 Chen, R. et al. A male-derived nonribosomal peptide pheromone controls female schistosome development. Cell 185, no. 9 (2022): 1506–1520 e1517, 10.1016/j.cell.2022.03.017

33 You, Y. et al. An improved medium for in vitro studies of female reproduction and oviposition in *Schistosoma japonicum*. Parasit Vectors 17, no. 1 (2024): 116, 10.1186/s13071-024-06191-y

34 Walker, A. J., Rinaldi, G. & Shakir, E. M. N. Molecular interactions between male and female schistosomes - a role for remote communication? Trends Parasitol 41, no. 1 (2025): 28–37, 10.1016/j.pt.2024.11.008

35 Du, P. et al. Proteomic and deep sequencing analysis of extracellular vesicles isolated from adult male and female *Schistosoma japonicum*. PLoS Negl Trop Dis 14, no. 9 (2020): e0008618, 10.1371/journal.pntd.0008618

36 Lu, Z. et al. Schistosome sex matters: a deep view into gonad-specific and pairing-dependent transcriptomes reveals a complex gender interplay. Sci Rep 6 (2016): 31150, 10.1038/srep31150

37 Lu, Z., Spanig, S., Weth, O. & Grevelding, C. G. Males, the wrongly neglected partners of the biologically unprecedented male-female interaction of Schistosomes. Front Genet 10 (2019): 796, 10.3389/fgene.2019.00796

38 Wendt, G. R. & Collins III, J. J. Schistosomiasis as a disease of stem cells. Curr Opin Genet Dev 40 (2016): 95–102, 10.1016/j.gde.2016.06.010

39 Korsunsky, I. et al. Fast, sensitive and accurate integration of single-cell data with Harmony. Nat Methods 16, no. 12 (2019): 1289–1296, 10.1038/s41592-019-0619-0

40 Agatsuma, T. Origin and evolution of *Schistosoma japonicum*. Parasitol Int 52, no. 4 (2003): 335–340, 10.1016/s1383-5769(03)00049-7

41 Shan, T. et al. Non-parasite genome encoded virus-like RNAs reprogram the pathogenicity of human blood flukes. Nat Commun 17, no. 1 (2025): 2995, 10.1038/s41467-025-67822-1

42 Howe, K. L., Bolt, B. J., Shafie, M., Kersey, P. & Berriman, M. WormBase ParaSite - a comprehensive resource for helminth genomics. Mol Biochem Parasitol 215 (2017): 2–10, 10.1016/j.molbiopara.2016.11.005

43 Love, M. I., Huber, W. & Anders, S. Moderated estimation of fold change and dispersion for RNA-seq data with DESeq2. Genome Biol 15, no. 12 (2014): 550, 10.1186/s13059-014-0550-8

44 Ou, Z. et al. Spatiotemporal transcriptomic profiling reveals the dynamic immunological landscape of alveolar echinococcosis. Adv Sci (Weinh) 12, no. 18 (2025): e2405914, 10.1002/advs.202405914

45 Cable, D. M. et al. Robust decomposition of cell type mixtures in spatial transcriptomics. Nat Biotechnol 40, no. 4 (2022): 517–526, 10.1038/s41587-021-00830-w

46 Cogswell, A. A., Collins III, J. J., Newmark, P. A. & Williams, D. L. Whole mount in situ hybridization methodology for *Schistosoma mansoni*. Mol Biochem Parasitol 178, no. 1-2 (2011): 46–50, 10.1016/j.molbiopara.2011.03.001

47 Pearson, B. J. et al. Formaldehyde-based whole-mount in situ hybridization method for planarians. Dev Dyn 238, no. 2 (2009): 443–450, 10.1002/dvdy.21849

48 Wang, X., Fang, C. & Cheng, G. Genome-wide identification and functional characterization of GPCR family genes reveal their key roles in the vitellarium development and egg production in *Schistosoma japonicum*. Parasit Vectors 18, no. 1 (2025): 286, 10.1186/s13071-025-06929-2

49 Livak, K. J. & Schmittgen, T. D. Analysis of relative gene expression data using real-time quantitative PCR and the 2(−Delta Delta C(T)) Method. Methods 25, no. 4 (2001): 402–408, 10.1006/meth.2001.1262

50 Wang, J., Chen, R. & Collins III, J. J. Systematically improved in vitro culture conditions reveal new insights into the reproductive biology of the human parasite *Schistosoma mansoni*. PLoS Biol 17, no. 5 (2019): e3000254, 10.1371/journal.pbio.3000254

51 Costain, A. H., MacDonald, A. S. & Smits, H. H. Schistosome egg migration: mechanisms, pathogenesis and host immune responses. Front Immunol 9 (2018): 3042, 10.3389/fimmu.2018.03042

52 Liu, J., et al. *Schistosoma japonicum* IAP and Teg20 safeguard tegumental integrity by inhibiting cellular apoptosis. PLoS Negl Trop Dis 12, no. 7 (2018): e0006654, 10.1371/journal.pntd.0006654

